# Nesting behaviour predicts heat tolerance evolution and climate vulnerability in bees

**DOI:** 10.1101/2025.09.02.673611

**Authors:** Carmen R. B. da Silva, Julian E. Beaman, James B. Dorey, Tessa Bradford, Tobias J. Smith, Ros Gloag, Vanessa Kellermann

**Affiliations:** School of Natural Sciences, Macquarie University, North Ryde, New South Wales, Australia; College of Science and Engineering, Flinders University, Bedford Park, South Australia, Australia; Environmental Futures Research Centre, School of Science, University of Wollongong, New South Wales, Australia; School of Biological Sciences, The University of Adelaide, Adelaide, Australia; School of the Environment, The University of Queensland, St Lucia, Queensland, Australia; School of Life and Environmental Sciences, The University of Sydney, Sydney, New South Wales, Australia; School of Agriculture, Biomedicine and Environment, La Trobe University, Bundoora, Victoria, Australia

**Keywords:** nesting ecology, behavioural thermoregulation, climate change, biogeography, phylogenetics, evolutionary history, physiological traits, heat tolerance, convergent evolution, repeated evolution

## Abstract

Species vulnerability to climate change depends in part on their capacity to evolve in response to increasing heat^1^. Within terrestrial ectotherms, heat tolerance generally corresponds weakly to current climates, which has led many to conclude that this trait is evolutionarily constrained^2–4^. However, most studies have not accounted for the role of microclimates, potentially obscuring signals of local adaptation. We examined heat tolerance in 95 species of wild bees that varied in nesting behaviour across the latitudinal extent of Australia. Species nest (ground, wooden cavities, or plant stems) micro-climate temperatures predicted heat tolerance evolution, where stem nesters evolved the highest heat tolerances, and ground nesters evolved the lowest heat tolerances due to their ability to evade extreme heat. A moderate level of phylogenetic inertia in heat tolerance was explained by patterns of related species sharing nesting behaviours. This indicated repeated adaptive evolution of similar heat tolerances, rather than strong evolutionary constraints on heat tolerance. Finally, incorporating nesting behaviour into assessments of climate change vulnerability changed the rank order of which species were most at risk. This underscores the need to understand what drives the evolution of heat tolerance across species to better identify the taxa most at risk to climate change.

## Main

Understanding and predicting the risks posed to biodiversity by climate change depend on understanding how tolerance to heat evolves^3–8^. A fundamental assumption is that an organism’s heat tolerance evolves in response to environmental heat exposure, such that biota in hot climates will be more heat tolerant than those living in cooler climates^4,9–11^. However, in terrestrial ectotherms the expected positive relationship between heat tolerance and environmental temperature is often weak or absent^2–4,12–14^. For example, species living at high and low latitudes often have similar heat tolerances^2,4,14,15^. Several explanations for this pattern have been proposed, including that heat tolerance reflects adaptation to past rather than present climates^5,12^, that heat tolerance in some taxa is evolutionarily constrained, due to low genetic variance or trade-offs^3^, or that species adapt to rare temperature extremes rather than climate averages, resulting in selection across geographic space that is more uniform than expected^12,16^.

Mismatches between species’ heat tolerances and their environments could also arise if macro-scale ambient temperatures are decoupled from the micro-climates in which animals actually live^17–19^. While macro-scale climatic variables are commonly used to assess associations between climate and heat tolerance, individuals experience their environment at finer spatio-temporal scales^19–21^. This is especially true for small animals such as insects and other arthropods. For example, species whose habitats are generally shaded, or that have partially subterranean lifestyles will be buffered against the ambient air temperatures of their macro-climate^17,18,22^, and evidence is emerging that thermal tolerances vary according to micro-climate in a range of animals^20,23^. Likewise, variation between species in behaviours that influence the micro-climates they typically experience, such as foraging or nesting behaviours, could impact selection on heat tolerance, and shape how heat tolerance evolves (i.e. the “Bogert effect” coined by Huey et al. 2003^22^). If so, assessments of the vulnerability of species to climate change may be improved by incorporating micro-climate^17,24^.

In this study, we investigated how heat tolerance evolves across 95 phylogenetically diverse species of adult bee (4 families, 22 genera, and 3,484 individuals) spanning the entire latitudinal gradient of mainland Australia (Figure 1A, ST1). Bees are keystone species in terrestrial ecosystems due to their role as pollinators^25^, including agro-ecosystems all over the world where they make vital contributions to crop fruit set, worth billions of dollars annually in the USA alone^26^. Bees are also a valuable system for understanding the scale at which climates shape thermal tolerance because co-occurring species may occupy different micro-climates due to differences in nesting behaviour. Some species nest in the ground, others in wood cavities (e.g. tree hollows or dead wood), while others nest in plant stems or existing holes in small twigs/branches (hereafter referred to as stem nesting bees)^25^. Based on these nesting behaviours, adults and young of stem nesting bees should be exposed to higher temperatures than bees that nest in wood cavities and the ground, due to the lack of insulation provided by thin stem nest walls. Similarly, bees that nest in the ground occupy the micro-climates most buffered from high air temperatures^27^. First, to determine the importance of nest micro-climate in dictating the evolution of heat tolerance we asked whether nesting behaviour can better explain inter-specific variation in heat tolerance than macro-scale temperature and precipitation data (i.e. regional weather station data that is used to model climatic variables at a 1km^2^ resolution), while also considering species’ evolutionary history (see native bee phylogeny SF1 and SF2). Next, we asked to what extent patterns in heat tolerance and nesting behaviour are consistent with either evolutionary constraint (phylogenetic inertia) or adaptation. We tested whether nesting behaviour has shaped the evolution of heat tolerance in bees independently of phylogeny by determining whether nesting behaviour influences the strength of selection, or mode of selection (i.e. whether heat tolerance evolves towards different optima depending on nesting behaviour), on heat tolerance. We then assessed phenotypic plasticity in heat tolerance in a thermally-variable population of bees to ensure our patterns in heat tolerance across nesting behaviours were not driven by plasticity. Finally, we compared climate change vulnerability across species with and without consideration of nesting behaviour, and demonstrated that including micro-climate drastically changes the rank order of species climate risk. With bees already threatened by warming and drying climates in some global regions^28,29^, we show that nesting behaviour can help identify those bees most at risk to climate change.

**Figure 1.**
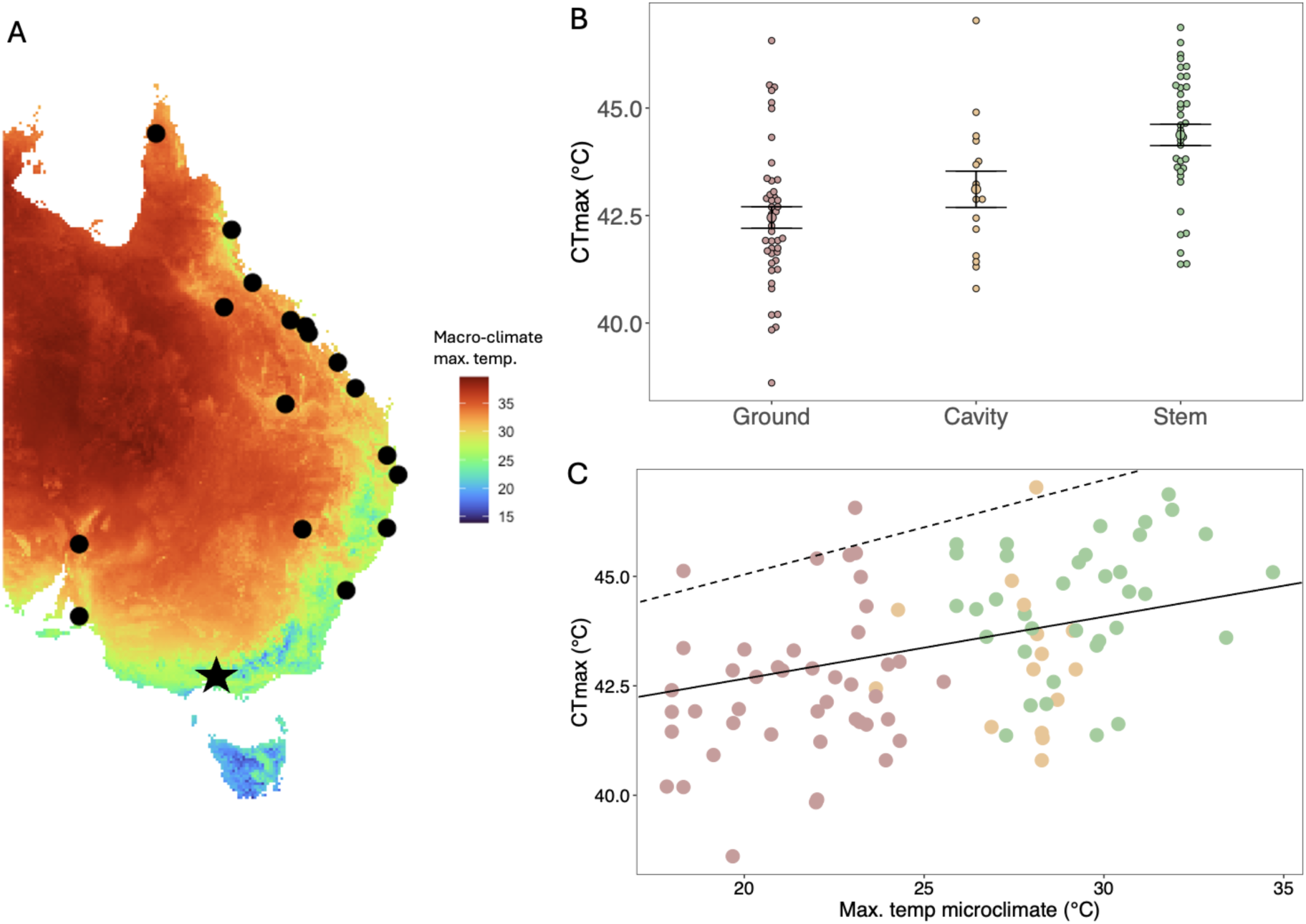
Native bee heat tolerance depends on their nest micro-climates. **A)** Sites at which native bees were sampled and assayed for heat tolerance. Star indicates the site at which intraspecific plasticity in heat tolerance was investigated. Map colours indicate macro-scale mean maximum temperature of the hottest month (BIO5^36^); **B)** Native bee heat tolerance (CT_MAX_) across nesting behaviours (mean and standard error bars; n = 95 species; **C)** Relationship between heat tolerance and micro-climate maximum temperature with species nesting behaviour indicated by colour; ground = brown, cavity = yellow, and stem = green. Solid line indicates the observed evolutionary regression slope and dashed line represents the optimal regression slope (i.e. unconstrained adaptive slope).

### Does heat tolerance evolve in response to the environment?

To first determine whether heat tolerance varied across species in a way consistent with adaptation to current climates we extracted macro-scale climate data (from the site of collection) for two key climate variables that are commonly used to assess the association between heat tolerance and climate: maximum temperature and mean annual precipitation^3,29,30^. Like many studies before us^12,31,32^, we found that these macro-scale climatic variables explained only a small proportion of between-species variation in heat tolerance (Ournstein-Uhlenbeck phylogenetic mixed model: R^2^ = 0.07, χ^2^ = 5.67, P = 0.017, model AICc comparison in ST 2 shows poor performance compared to other models). Macro-scale estimates of climate, however, are unlikely to capture the temperatures that small ectotherms, such as bees, experience in their nests^19,20^. To capture nest micro-climates we extracted the predicted maximum temperatures and mean humidities species experience within ground, wood cavity, or stem nests (see details in Materials and Methods). Maximum nest micro-climate temperature, but not humidity, explained more variation in heat tolerance than macro-scale climate alone (Ornstein-Uhlenbeck phylogenetic mixed model: R^2^ = 0.12)(ST 2), such that, on average, bees nesting in stems across the latitudinal gradient were the most heat tolerant and those nesting in the ground were the least heat tolerant (Likelihood ratio test (LRT): χ^2^ = 11.82, P < 0.005) (Figure 1B & C & Figure 2) (ST 2). This pattern suggests that ground-nesting behaviour buffers bees from selection for high heat tolerance, while stem nesting bees are adapted to their hotter micro-climates (Figure 1B & 1C; Figure 2) i.e. the Bogert effect^17,22^. Plotting the residuals of the relationship between heat tolerance and macro-scale temperatures across the phylogeny further supported that ground nesting bees are buffered from selection, as they had lower than expected heat tolerances given the macro-scale temperatures they experience, while stem nesting bees showed the opposite pattern (Figure 2). However, whether this pattern reflects weaker selection for high heat tolerance at the adult life stage in ground nesting bees (as they use behaviour to escape hot temperatures throughout the day) or instead reflects selection towards lower heat tolerances at pre-adult stages, is unknown. We also examined whether body size was an important predictor of heat tolerance, given that smaller species exchange heat more rapidly than larger species^33^. However, we found no relationship between body size and heat tolerance (LRT: χ^2^ = 2.01, *p* = 0.156) (ST2). This result contrasts some recent studies that have found heat tolerance is related to body size in insects, including bees^29,34^, but is in line with studies in *Drosophila*^35^. We also found no association between body size and latitude (i.e. no support for Bergmann’s rule) (LRT: χ^2^ = 1.85, *p* = 0.174) (SF 3 & 4).

**Figure 2.**
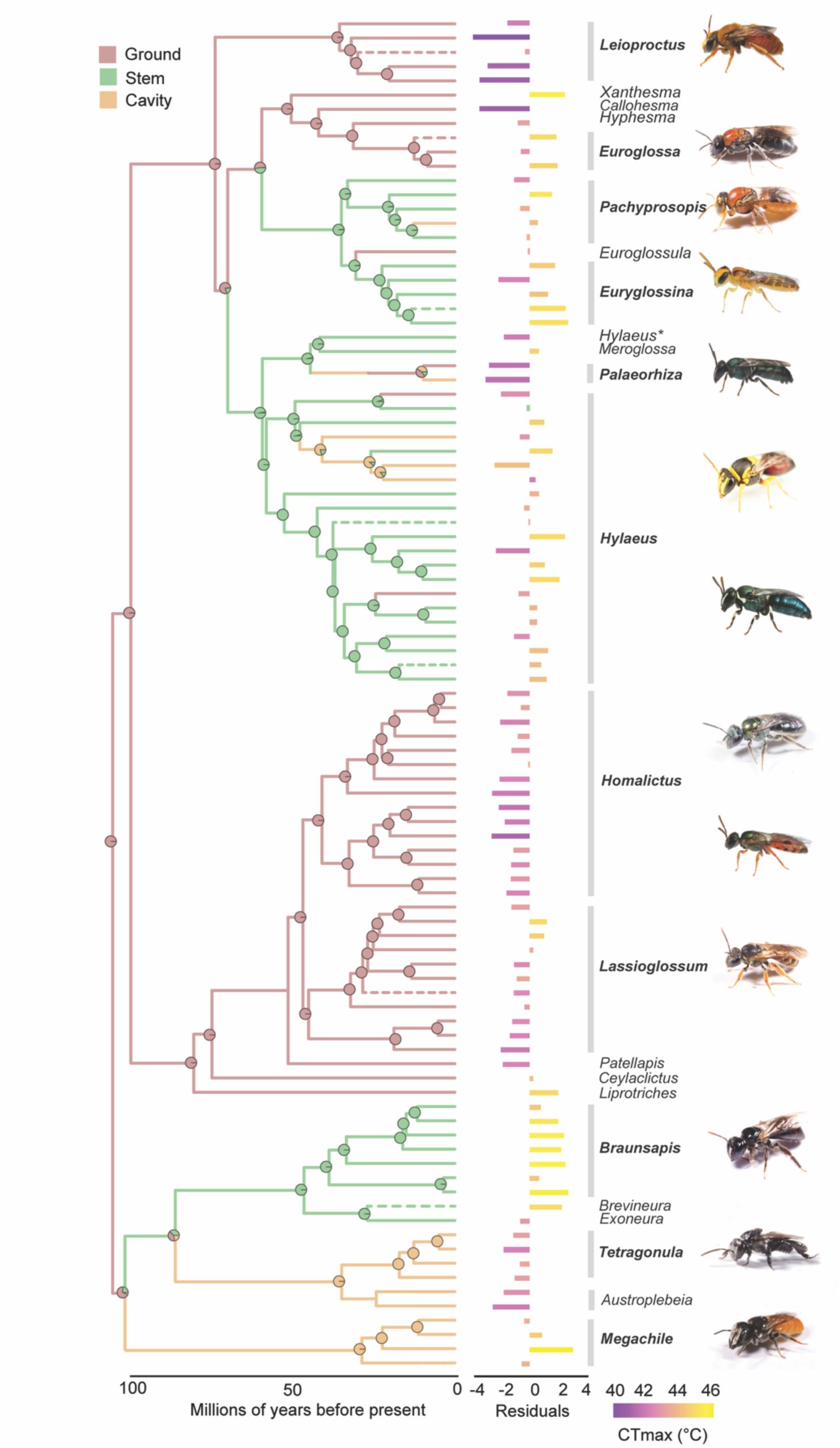
Nesting behaviour and heat tolerance (CT_MAX_) of native bees. Pie charts represent the posterior probabilities of nesting ancestral states at each internal node. Bar plots represent the heat tolerance residuals from an OU phylogenetic mixed model and indicate whether bees have higher or lower thermal tolerances relative to their mean thermal macro-climate (BIO^36^); ground nesters tend to have low thermal tolerances for their macro-climate and stem nesters tend to have high thermal tolerances for their macro-climate (bar colour indicates heat tolerance). The final column represents the genus of each species tested. Bee photos were provided by JD, TS and Nick Volpe. *note: *Hylaeus* appears twice in the phylogeny. This is because our phylogeny suggests that *Hylaeus nubilosis* actually sits in the clade containing *Paleaorhiza* and *Meroglossa* (SF 1), where they have been noted to have similar facial features^37^.

Studies in ectotherms have found strong signals of evolutionary history in heat tolerance, which have often been inferred to mean there are constraints in the capacity to evolve heat tolerance^3,5^. For example, some lineages might lack the standing genetic variance required to adapt rapidly with changing environments, resulting in a strong phylogenetic signal and heat tolerances that are poorly matched to their current environments^38^. However, phylogenetic signal in a trait does not necessarily imply constraints and could arise from adaptation^39^. If closely related species occupy and adapt to similar environments (e.g. similar nest micro-climates), these species will have similar trait values. Phylogenetic signal in this context more likely reflects phylogenetically structured adaptation.

To examine whether there is evidence of phylogenetically structured adaptation, we first looked at whether phylogenetic signal was present in the heat tolerances of bees by examining the phylogenetic half-life (*t*_1/2_) of an intercept-only Ornstein-Uhlenbeck process model^3, 39^. The phylogenetic half-life represents the average time it takes (in the units of the phylogeny) for a trait to evolve halfway from its ancestral state towards a new optimum^39^. A low *t*_1/2_ ratio (*t*_1/2_ ratio = phylogenetic half-life divided by tree hight) indicates fast adaptation, a high *t*_1/2_ ratio reflects slow adaptation (strong phylogenetic inertia), and a *t*_1/2_ ratio over 1 (i.e. *t*_1/2_ greater than the height of the tree) represents evolution via Brownian Motion. We found moderate phylogenetic signal, suggesting that heat tolerance was not evolving independently from the phylogeny, nor by Brownian Motion (*t*_1/2_ = 35.79 MY; tree height = 104.12 MY; *t*_1/2_ ratio = 0.34, CI = 0.22-0.47; SF 3). We then compared this to the phylogenetic signal in the Ornstein-Uhlenbeck process model that examined the relationship between heat tolerance and micro-climate temperature (ST2) to tease apart whether the phylogenetic inertia represents phylogenetically structured adaptation or constraints. The estimated *t*_1/2_ decreased (*t*_1/2_ = 26.64 MY; *t*_1/2_ ratio = 0.25, CI = 0.16 – 0.35), indicating that some of the phylogenetic signal in heat tolerance is explained by adaptation to micro-climate temperature. The remaining ∼25% of the phylogenetic signal from the intercept only model reflects low to moderate phylogenetic inertia^39^. Thus, heat tolerance is partly constrained by shared ancestry, but species have also evolved higher heat tolerances with warmer micro-climate temperatures (evidenced by the difference between the evolutionary and the optimal regression slopes in Figure 1C) (see ST 3 for full model coefficients).

### How does nesting behaviour influence heat tolerance evolution?

Another approach to understanding how nesting behaviour (and thus micro-climate temperature) influences the evolution of heat tolerance is to consider similarities between lineages in which a given nest type has evolved independently. Nesting behaviour is phylogenetically structured, such that only two independent transitions from the ancestral state of ground nesting to both stem and ground nesting have occurred deep within the phylogeny giving rise to the stem/cavity nesting species we see today (Figure 2, ST 4). Within the stem and cavity nesting species, five and two transitions have occurred back to ground nesting, respectively (Figure 2, ST 4). The strong association between nesting behaviour and phylogeny means it is likely that nesting behaviour has influenced the strength or mode^40^ of selection on heat tolerance. If ground nesting bees experience weaker selection on high heat tolerance because they have greater potential for behavioural thermoregulation, and thus evolve more slowly towards their thermal optima than bees in more exposed nest types, then we expected to detect differences in the rate heat tolerance evolves depending on nesting behaviour. If bees with different nesting behaviours evolve at similar rates but towards distinct thermal optima, then we expected to detect differences in the mode of selection according to nesting behaviour. Such differences in selection mode would provide additional evidence that bees adapt to the temperatures of their nest micro-climate. Furthermore, the strong phylogenetic structuring of nesting behaviour could also explain the observed phylogenetic inertia in heat tolerance (above); that is, closely related species may share similar heat tolerances because they experience the same selective pressures. To better tease apart the manner in which nesting behaviour has influenced heat tolerance evolution we used two complementary analyses.

First, we modelled how heat tolerance varies across the phylogeny to ask whether nesting behaviour principally influences the strength (rates) or mode of selection (i.e. optima at which thermal tolerance evolves towards). We compared a number of different models contrasting single and multiple rates and optima (for a comparison of models see Methods and ST 5) and found that heat tolerance was evolving towards distinct optima for each nesting behaviour, but that the rates of evolution were the same across nesting behaviours (σ^2^ = 0.34°C per MY, low 95 CI = 0.30–0.36) (lowest AICc, ST 5). Stem nesters had the highest phenotypic optima (**ϑ** = 44.90°C, 95 CI = 44.06–45.43), followed by cavity nesters (**ϑ** = 43.27°C, 95 CI = 43.27–43.93), and ground nesters (**ϑ** = 42.88°C, 95 CI = 42.88–43.45) (Figure 2) (ST 6). That is, nesting micro-climates impact the mode of heat tolerance evolution. While we did not explicitly test a multi-rate and multi-optima model (because these models are known to be unreliable when working with less than a few hundred species^41^), if nesting behaviour influenced both the strength and mode of selection we expected that both the multi-rate and multi-optima models would perform similarly, however the multi-optima model considerably out-performed the multi-rate model (ST 5). Moreover, these models show that phylogenetic inertia alone does not explain heat tolerance, as we would then have expected support for a single rate model with no evolutionary optima. Instead, distinct selective modes are consistent with repeated evolution of heat tolerance among bees with shared nesting behaviours, suggesting adaptation to nest micro-climate underpins the evolution of heat tolerance.

Our second approach assessed whether heat tolerance has evolved repeatedly depending on nesting behaviour. Using the Wheatsheaf index (*w*) we measured the strength of repeated evolution while accounting for phylogenetic relatedness. We also tested whether species with the same nesting behaviour had more similar heat tolerances than expected given that nesting behaviour is highly structured across the phylogeny (*P*). We again found evidence of repeated evolution of heat tolerance across all nesting behaviours. That is, species with similar nesting behaviours had similar heat tolerances irrespective of phylogeny. This was particularly true for both stem and ground nesters (Table 1. P < 0.05), with a weaker signal of repeated evolution in cavity-nesters, possibly due to the lower sample size for this group. In all, our estimates of distinct phylogenetic optima and repeated evolution in heat tolerance for bees with different nesting behaviours point to adaptive processes driving heat tolerance evolution. Indeed, this is the first example of repeated evolution in heat tolerance associated with a common behaviour.

**Table 1.** Wheatsheaf index (a measure of repeated evolution) and upper and lower 95% confidence intervals. The P-value tests whether the evidence of repeated evolution is particularly strong given the topology of the phylogenetic tree^42^.

| Nesting behaviour | w | Lower CI | Upper CI | P-value |
| --- | --- | --- | --- | --- |
| Stem | 0.97 | 0.95 | 1.02 | 0.016* |
| Cavity | 1.26 | 1.23 | 1.30 | 0.305 |
| Ground | 0.84 | 0.83 | 0.86 | 0.032* |

### Does phenotypic plasticity or population variation explain trends in heat tolerance?

An alternative explanation for why heat tolerance varies with nesting behaviour is that bees in stem nests could be acclimated to higher temperatures than bees nesting in the ground. That is, phenotypic plasticity could underpin differences in heat tolerance between stem, cavity, and ground nesters. To test whether phenotypic plasticity was contributing to differences between our stem and ground nesting bees, we sampled and assayed bees for heat tolerance once a month across the active foraging season at our most climatically variable high-latitude site (Melbourne, Fig 1A; Nov-March, 542 individual bees assessed, 26 species with 3–80 individuals per species). This site was chosen because bees at higher latitudes experience greater seasonal variability in temperature and thus are the most likely to show plasticity in thermal tolerances based on the climate variability hypothesis (phenotypic plasticity increases with increased environmental variability^43^) (Figure 1). We predicted that if phenotypic plasticity was driving the difference between stem and ground nesting bees then stem nesters at our high-latitude site would exhibit greater seasonal plasticity than sympatric ground nesters as the former are exposed to greater thermal variability. However, heat tolerance did not differ across the months of collection for either ground nesters or stem nesters (Fig. 3A & B) (LRT: χ^2^ = 4.10, *p* = 0.392). For nine species of bees, where we had estimated heat tolerance in two or more months, we looked at whether variation in heat tolerance could be explained by species (40.94% of variation explained) or by month/season (9.61% of variation explained) (Figure 3C) (ST 7). The much higher proportion of variation being partitioned to the species level indicates limited seasonal plasticity in heat tolerance. Therefore, plastic differences in heat tolerance are unlikely to explain the differences we observe between stem, cavity and ground nesting bees. Plasticity could also underpin the significant relationship between heat tolerance and micro-climate temperature (Figure 1C), but given the limited evidence for heat tolerance plasticity in our highest latitude populations of bees we think this is unlikely. More generally, little or no plasticity in heat tolerance within bee species is consistent with a growing body of literature across ectotherms^44^.

**Figure 3.**
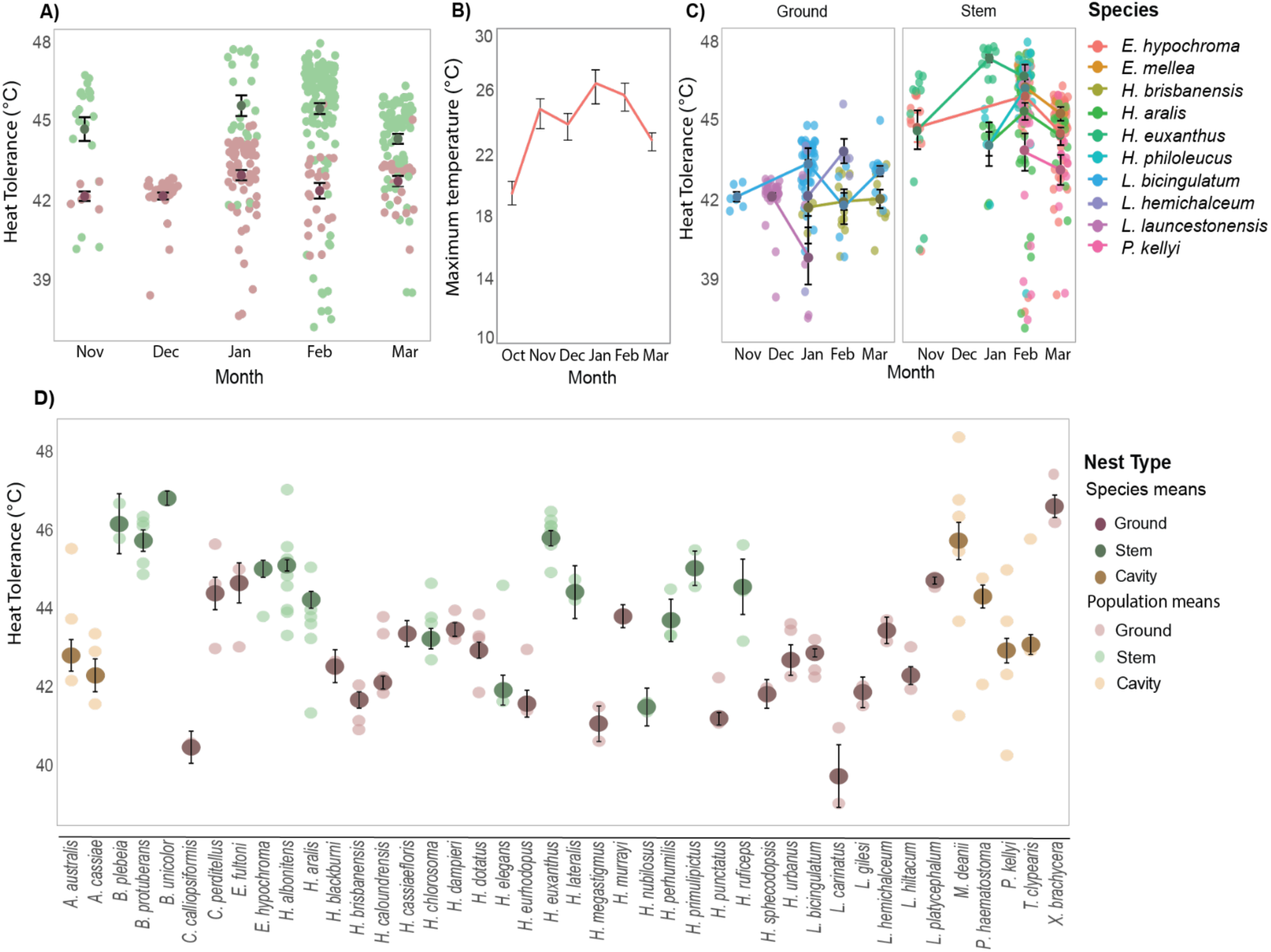
**A**) Heat tolerance across the active period of bees (November – March) in Melbourne. Each data point reflects individual estimates, with stem-nesters in green and ground-nesters in brown **B)** Seasonality in maximum temperature at this sample site. Note October was included as the month prior to trait estimation is likely to influence the plastic response. **C)** Heat tolerances for ten species of bees where we estimated heat tolerance for at least two months across the active period of bees (November – March). Individual data points and means for each species (+/-one standard error) D) Population means and species means (+/-one standard error) for heat tolerance for 43 species of bees.

Finally, given that only a single population was sampled for many of the 95 species in our total dataset, it is possible that the sampled population is not reflective of the species’ overall heat tolerance (i.e. different populations within a species might show local adaptation in heat tolerance). To assess whether local adaptation might have influenced our findings, we partitioned heat tolerance variation at the species and population level for 43 species in which we had sampled at least two populations. Here we found that 29.15% of the variation in heat tolerance was attributable to species, while 7.37% of the variation was attributable to populations (63.48% residual variance). That is, variation between species was greater than the variation within species (Figure 3D). Therefore using only a single (or averaged) population estimate is unlikely to have confounded the broad trends that we observed between species with different nest types (Figure 3D).

### Does micro-climate influence predictions of species vulnerability?

A major goal in evolutionary ecology is to predict species vulnerability to climate change^1,6^. Common metrics used in these predictions include thermal safety margins and estimates of species range shifts with niche models^6,32,45^. The micro-climates organisms occupy are rarely factored into vulnerability predictions (but see^17,24^). To test the importance of nesting behaviour (which determines bee micro-climate) on climate change vulnerability, we estimated thermal safety margins as the degrees distance between species heat tolerance and the maximum temperature of their environment.

To emphasise the importance of how behaviour can influence species micro-climates, we first calculated species thermal safety margins based on the air temperatures bees were collected from. Here we found that ground nesters tended to be the most vulnerable, due to their lower thermal tolerances (Figure 4A). However, when we calculated thermal safety margins considering nesting micro-climate temperature, we found that this trend in vulnerability was reversed (Figure 4B & C). That is, species nesting above ground (in stems and cavities) had the narrowest safety margins (Fig 4B & C). This is because the higher heat tolerances of stem and cavity nesters do not compensate in full for the hotter temperatures to which they are exposed (Figure 1C, SF3). This pattern is consistent across geographic space, though margins become narrower for all species at locations closer to the equator (Figure 4B). In other words, for any given location, stem nesters have narrower safety margins than sympatric ground nesters. Our analysis reveals that, in the absence of data on heat tolerance, species nesting behaviour could be a suitable proxy for ranking species vulnerability in a given location. We also show generally high heat tolerances in bees and subsequently broad thermal safety margins, similar to other diurnal insect pollinators such as butterflies^15^. But sublethal effects of temperature are likely to impact populations at much lower temperatures than those calculated by safety margins^47,48^, hence safety margins should be used to rank species vulnerabilities, not predict the temperature at which they will go extinct^46^. Finally, we only considered adult bees, thermal safety margins of pre-adult stages could differ.

**Figure 4.**
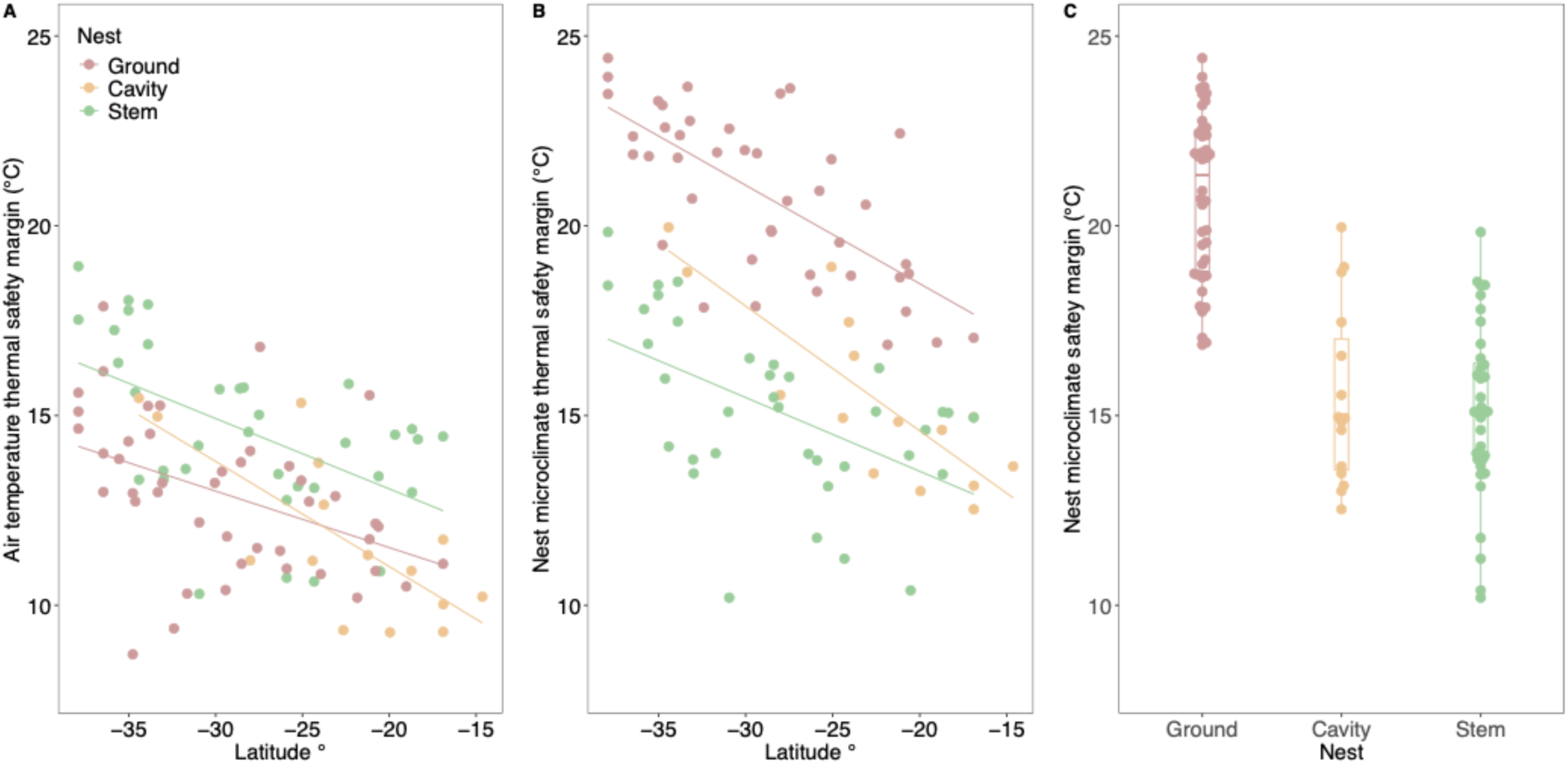
Thermal safety margins of bees depending on. **(A)** species air temperature thermal safety margins and, **B)** nest micro-climate temperature thermal safety margins, across latitude, and **C)** Nest micro-climate thermal safety margins.

Our results suggest stem nesters have the lowest capacity to escape unfavourable environmental temperatures and are likely to be the most impacted by anthropogenic climate change in the near term. Importantly though, as all bees must forage outside their nests and therefore be exposed to a shared climate when foraging, even ground-nesters will be impacted from warming climates in the longer term. Indeed, some models predict that thermoregulators (in this case, ground-nesting bees) will be the least at risk in the short term, but the most at risk in the long term, because the buffering effect of behaviour on the evolution of heat tolerance will drive an increasing mismatch between the heat tolerance that they will need to survive climate change. Near-term risks, however, are those most relevant to today’s population management and conservation decisions. Given we also show that phenotypic plasticity is unlikely substantially increases heat tolerance in the short term, which is an emerging them in ectotherms in general^44^, our estimates of safety margins are likely to capture near term risk. Our results highlight the importance of factoring in micro-climates based on differences in species behaviour into predictions of climate change vulnerability (Figure 4 A & B).

## Conclusion

Here we show that associations between adult bee heat tolerance and thermal environment are stronger when we consider a key behaviour that determines species’ micro-climates. By accounting for nesting behaviour, bees in our study therefore adhere to the expected, but rarely found, relationship between heat tolerance and environmental temperature^2,4,12^. Selection on heat tolerance acts at the level of the nest micro-climate, such that bees that nest in the ground evolve towards different evolutionary optima in heat tolerance than those that nest above ground in stems or tree cavities. Although outside the scope of this study, other behaviours that may modify the nest micro-climate might similarly shape how heat tolerance evolves (e.g. sociality^50^). The implications for predicting the vulnerability of bees and other ectotherms to changing climates is two-fold. First, confirmation that heat tolerance evolves with increasing environmental temperature supports the view that bees may also adapt to future climates. Whether such evolutionary responses will be enough to keep pace with changing climates remains an open question, but even small adaptive shifts in heat tolerance can substantially reduce extinction risk^51^. Second, even where future evolutionary responses are set aside, understanding what drives the evolution of heat tolerance across species can help to identify at-risk taxa. In bees, stem nesters have evolved the highest heat tolerances but also occupy the warmest micro-climates and thus have the narrowest thermal safety margins. Including the relevant micro-climates that species occupy into models that predict and rank species vulnerability to climate change may improve our ability to determine the winners and losers of climate change.

## Materials and Methods

### Bee collection

We collected bees from 16 sites on the eastern side of Australia during two field trips. The first field trip took place January – March 2020 and the second October 2020 – February 2021 (see supplementary ST8 collection sites and order of collection). Bees were collected from flowering plants by sweep netting and were placed into 30 mL collection vials with foam plugs to allow airflow. We then placed vials of bees into a portable cooler with ice bricks that have been cooled in the refrigerator (wrapped in tea towels) to maintain bees at a comfortable temperature. We provided bees with a piece of paper towel dipped in 40% sugar solution to maintain homeostasis prior to thermal assays.

### Heat tolerance assays

We estimated the heat tolerance of individual bees following the protocol of da Silva et al 2021^27^. Briefly, each bee was placed into a watertight glass 5 mL testing vial and vials were labelled 1–150. Bee vials were placed haphazardly into two 75-vial testing racks. Once loaded, we placed racks into a water bath set to 26 °C. Water temperature was then increased by 0.1°C per minute using a TeTechnology^Ⓡ^ LC 200 liquid heater/cooler, controlled by an TETechnology^Ⓡ^ TC 720 temperature control device and powered by a Peltier power supply. Water was circulated using a pond pump and the water temperature was increased until each individual bee lost complete movement. The time and temperature that each bee reached their critical upper thermal limit (CT_MAX_) was recorded. At the end of each assay, we placed bees into 99% ethanol for later identification and genotyping.

### Wing length

Because body size is associated with thermal tolerance in some studies^34,52^, we assessed whether it explains significant variation in heat tolerance, using wing length as a proxy for body mass as they are tightly correlated^53^. We removed the right wing from 10 individuals haphazardly selected from each species (or as many individuals as possible depending on the number of individuals tested per species, sample sizes provided in Supplementary Data). Wings were mounted on microscope slides in a solution of ethanol and glycol. A photograph of each wing was taken using a Lecia M80 stereo microscope (Leica, Heerbrugg, Switzerland). Wing length was measured from where the wing attaches to the thorax, to where the upper wing vein ends (SF 4), using ImageJ.

### Bee Identification

Bees were first identified to family, genus and subgenus using keys in Smith (2018)^54^ and Michener (2007)^25^. Species-level identifications were subsequently undertaken using a range of keys:^25,54–74^. Keys were used in conjunction with the online photo libraries and digital keys on the PaDIL Pollinators website (www.padil.gov.au/search/pollinators), digital keys by Michael Batley, Australian Museum (https://michaelbatley.github.io/Bee-ID-SH/main.htm, https://michaelbatley.github.io/Bee-ID-SM/main.htm) and online distribution maps (Atlas of Living Australia, www.ala.org.au and iNaturalist, www.inaturalist.org). In several cases, identifications of species that lack keys were made with assistance from bee taxonomists at the Australian Museum and Museums Victoria (see Acknowledgements). Voucher specimens were lodged in the Kellerman Lab reference collection, La Trobe University.

For bees that could not be confidently identified using taxonomic keys, we instead used mitochondrial CO1 barcoding to generate a species ID. As many Australian bee species have not yet been formally described, those that did not match a specimen on BOLD or GenBank were assigned a morpho-species name (see sequencing details below).

### Nesting behaviour

We assigned bees to one of three nesting categories (ground, stem, or cavity nesting) based on nest type descriptions in taxonomic and reference texts (cited in Supplementary data). The category stem includes bees living in plant stems, twigs, and small narrow branches as they have similar thermal properties. Some species, such as those in the genus *Hylaeus,* can sometimes be found nesting in stems, twigs, or existing cavities in larger branches or trunks. For these species we allocated nesting strategies to the category where there were more numerous reports in the literature. A summary of each species physiological, climatic and nesting behaviour can be found in ST 1.

### Ultraconserved Element Phylogeny

We selected 75 species from our total dataset for ultraconserved element sequencing. Species with high sample sizes and those that represented unique subgenera within the collection were prioritised. DNA was extracted at the South Australian Museum from a single leg using a modified version of the Canadian Centre for DNA Barcoding Glass Fiber Plate DNA Extraction Protocol in PALL Acroprep ADVANCE 96-Well Filter Plates (1 ml 3.0 µm glass fibre^75^). The Meyer and Kircher (2010)^76^ protocol was followed to prepare indexed libraries for Illumina sequencing, incorporating the on-bead method of^77^ and^78^. Indexed samples were quantified fluorometrically on a Quantus fluorometer (Promega, Madison,WI, USA) and pooled into groups of five for the hybridisation step, based on similar concentration. MyBaits UCE Hymenoptera 2.5Kv2AB kit (Arbor Biosciences, USA) was used to capture UCE sequences, following the myBaits User Manual version 5.01. DNA concentration of the capture reactions was measured on a Quantus fluorometer and the libraries were pooled in equimolar quantities. Final fragment length distribution and molarity of the bee UCE library was determined on a TapeStation 2200 using high-sensitivity ScreenTape and reagents (Agilent Technologies, Santa Clara, CA, USA). Sequencing was conducted on a NovaSeq 6000 SP single lane at the Australian Genomic Research Facility (AGRF) in Melbourne with 150 bp paired-end read cycles.

We used the program Phyluce version 1.7.1^79^ to assemble the UCE contigs, check assemblies, match probes, generate fasta files, and align sequences. We used IQ-TREE 2 version 2.2.0^80^ for UCE phylogenetics with modelFinder^81^ using 1,000 bootstraps, and MF models. Using the scheme created by modelFinder, we employed IQ-TREE to build a constrained tree with the Apoid wasp outgroups *Ammophila cybele, Oxybelus analis*, and *Sphecius hogardii*. Original UCE phylogeny is shown in SF 2.

We used Timetree^82^ to generate median divergence times for each of the family clades based on previous phylogenetic studies^83–92^.

### Adding further branch tips via CO1 sequencing

In addition to the specimens chosen for UCE sequencing, our heat tolerance dataset included some additional species (for which >5 individuals were assayed). CO1 data was collected from GenBank and BOLD (https://boldsystems.org/) for 34 species (some species for which we had no UCE data, and some species so that we had an overlap in UCE and CO1 data to make an integrated UCE and CO1 tree. For the species where no CO1 data was available online, we sequenced the remaining 29 species to ensure that these additional species could be included in analyses. We extracted DNA from one specimen per species using a 5% chelex solution. We then amplified a ∼655 bp region of CO1 (primers: LCO1490 and HCO2198^93^) and performed Sanger sequencing (Macrogen Inc, Seoul). COI sequences were then trimmed and cleaned for ambiguous base calls, and species IDs checked via BLAST and BOLD (https://boldsystems.org/). We then generated separate CO1 phylogenies for each genus using IQ-TREE.

To integrate species from the CO1 phylogenies into the UCE phylogeny, we standardized CO1 tree branch lengths to match the UCE tree using a scaling factor. This factor was calculated for shared species between the two trees. The scaling factor was derived by summing branch lengths from the genus or sub-genus root in both trees, dividing the CO1 sum by the UCE sum, and averaging this ratio across all shared species when multiple common species were present. Finally, we divided the CO1 branch lengths by the scaling factor to align them with the UCE tree. The final phylogeny used in our analysis included 52 UCE samples (not all UCE sequences were included in the trait analysis as those species had low sample sizes) which made up the backbone of our phylogeny (at least one UCE sample was included for each genus in the combined phylogeny) and 43 CO1 tips (N = 95).

### Climate and micro-climate data acquisition

We extracted historical climate data (1970–2000) for each collection site from WorldClim2^36^. We extracted maximum temperature of the hottest month (BIO5) and mean annual precipitation (BIO12) at a 30 arc seconds resolution using the R package *raster*^94^. We then calculated the mean hottest temperature of the hottest month and mean annual precipitation for each species across their collection locations to use in our phylogenetic mixed models assessing whether heat tolerance is correlated with climatic factors. We also extracted micro-climate data for each species depending on their nesting substrate using *NichmapR v3.3.2*^95^ and a global surface climate database^96^. For ground nesting species, we extracted maximum soil temperature at a depth of 100 cm in full sun (many *Lasioglossum* and *Homalictus* species which dominate our ground nesting species nest at this depth^37^). For stem nesting bees we collected maximum air temperature at 1.2 m above-ground because thin stems are unlikely to provide substantial thermal buffering for bees. Because cavity nesters are likely to be somewhat buffered from extreme temperatures (e.g. stingless bees nesting in tree hollows), but less so than bees that nest deep in the soil, we took the mean temperature of ground (100 cm in soil) and stem (1.2 m in the air) habitats and applied this to the cavity nesters at each collection site. We calculated micro-climate humidity for each collection site/nest type combination using the same methods as above; however, as cavity humidity is likely to more closely resemble air humidity than soil humidity,we therefore estimated cavity humidity as 75% of air humidity at the same site. As for regional climate variables, we used the mean maximum nest micro-climate temperature and humidity experienced by a species across all of their collection sites.

### Phylogenetic trait analyses

Phylogenetic trait analyses were conducted in the program *R v4.5.0*. While most phylogenetic comparative methods control for phylogenetic non-independence assuming traits evolve according to Brownian motion (BM; random movement), Ornstein Uhlenbeck (OU; evolution towards an optimum) process models more accurately describe trait evolution as they are able to treat trait evolution as a stochastic process (e.g. genetic drift) and a deterministic process (e.g. evolution towards phenotypic optima). Phylogenetic comparative methods that use an OU process, which can be used to examine evolution towards trait optima and control for phylogenetic inertia, have been argued to be more suitable for testing evolutionary hypotheses using physiological trait data^39,97,98^. In the case where traits do not evolve towards an optima, models revert to using a BM model of evolution^97,98^. Thus, we used the package *slouch v2.1.5*^99^ to determine how phylogeny, local climate (maximum temperature of the hottest month (BIO5) and annual precipitation (BIO12), and micro-climate (estimates of maximum temperature and mean humidity within the nest) shape heat evolution.

We computed models based on different predictions we had for how CT_MAX_ might evolve across bee species (ST 9). We asked whether macro-climate variables or micro-climate variables better explained variation in heat tolerance across the cline (local climate and micro-climate variables were not mixed within models). We conducted model comparison using AICc (Akaike Information Criterion) to determine which model best explains variation in heat tolerance evolution. We also examined the relationship between micro-climate temperature and latitude (SF 7). We put wing length into the best fitting model and assessed whether it explained significant variation in heat tolerance. We conducted a grid search to assess for measurement error. We assessed the significance of the factors in the best fitting model by conducting likelihood ratio tests.

### How does nesting behaviour drive heat tolerance evolution?

To determine how nesting behaviour shapes the rate or mode of heat tolerance evolution, we simulated nesting behaviour across the native bee tree 1000 times using the make.simmap function from the phytools *v2.4-4* package^100^. To determine the number of nesting behaviour state changes across the phylogeny (and the confidence in each node state) we calculated the posterior probabilities of each internal node. We then computed and compared a range of OUwie models^101^ using the 1000 character mapped phylogenetic trees as per^98^ (Figure 2). We compared evolutionary models where heat tolerance evolves at a single evolutionary rate (BM1); different evolutionary rates depend on nesting behaviour (BMS); evolves towards a single trait optimum (OU1); or evolves towards different trait optima depending on nesting behaviour (OUM). These analyses were iterated across the 1000 trees for the BMS and OUM models, but not for the BM1 or the OU1 models as they do not depend on simulated ancestral states. Models were compared using AICc (Akaike Information Criterion). We assessed the robustness parameter estimates, and whether there was likely to be significant differences between CT_MAX_ evolution depending on nesting behaviour by conducting a bootstrap analysis with 1000 simulations as per^98^.

To investigate whether patterns in heat tolerance are driven by nesting behaviour and not solely shared ancestry we calculated the wheatsheaf index (*w*), which is a measure of repeated evolution. It measures and compares the euclidean distance between species traits while considering relatedness^102^. We estimated *w* for each nesting behaviour, and assessed whether heat tolerance is more similar within each of these groups than we would expect at random by conducting a bootstrap analysis using the package *windex 2.1.0* ^42^.

Finally, we examined how one type of behaviour (i.e. differences in nesting substrate) influences the evolution of heat tolerance but this ignores other types of behavioural syndromes (i.e. site fidelity, timing of foraging activity, evaporative cooling). In bees, sociality (solitary, communal, eusocial) might be important^51^, as some social bees are known to thermoregulate their nests^103^. Australia has only three genera of eusocial bees (the stingless bees *Tetragonula* and *Austroplebeia and Exoneurella tridentata*) and the social status of many other Australian bee species however is unknown or variable. For example, in some populations of *Braunsapis* and *Exoneura*, some nests have a solitary female, while others have multiple egg-laying females^104,105^. Thus, in this study we were unable to include sociality into our analysis. However, as more data on Australian bee sociality becomes available in the future, they might help to reveal how social behaviours contribute to heat tolerance evolution.

### Phenotypic plasticity

To determine whether seasonal plasticity contributed to differences in CT_MAX_ within our dataset, we examined how heat tolerance varied among species across their active period during the summer months in Melbourne. We chose Melbourne because it is the most seasonally variable site, and we would therefore expect that if plasticity was contributing significantly to the variation in our results we would observe the biggest plastic shifts in CT_MAX_ in Melbourne across the season. This does not rule out the possibility that developmental plasticity contributes to plasticity in CT_MAX_. Bees were collected via sweep netting over native plants once a month between November 2020 to March 2021. Each month between 70 and 120 bees were assessed for CT_MAX_ using the methods above.

We looked at seasonal variation in heat tolerance using two approaches. The first approach included all individuals assessed for heat tolerance (27 species with a minimum of three individuals, Fig 3A). We used a mixed effects linear model in the package *lme4 1.1-37* ^106^ in R to ask whether heat tolerance varied across month and nest type, with month and nest type deemed fixed effects and species as a random effect. This analysis allowed us to ask whether heat tolerance changed across months and whether species with different nesting ecologies differed in how their heat tolerance changed across the months, which we would interpret from testing the interaction between month and nest type. This analysis did not allow us to explicitly ask whether individual species varied in their heat tolerance across months, which would require individual species to be sampled across multiple seasons. For our second approach we created a second dataset that included only species with at least three individuals per month, assessed in at least two months (10 species, Fig 3B). Using this dataset as above we used a mixed effects linear model to ask whether heat tolerance varied across month and nest type, with month and nest type deemed fixed effects and species as a random effect. For this dataset we were also interested in whether variation in heat tolerance could be portioned to the species level or the month. We quantified the contribution of different sources of variation by fitting linear mixed-effects models and partitioning total phenotypic variance into components attributed to the random effects species and month and the residual variation.

### Population comparison

To determine the extent to which populations varied in their heat tolerance, we reduced the full bee dataset to species where we had sampled more than three individuals from at least two populations (43 species). Using these data we quantified the contribution of different sources of variation (population/species) by fitting linear mixed-effects models and partitioning total phenotypic variance into components attributed to the random effects species and population and the residual variation.

### Thermal safety margins

To rank species vulnerability to climate change, we calculated each species thermal safety margin as the degrees distance between their heat tolerance and their nesting micro-climate temperature. We then compared these to thermal safety margins that were calculated using air temperatures only, as is the standard method^32^. This comparison was made to emphasise the importance of considering individual species behaviour when calculating risk to climate change.

## Acknowledgements

We would like to thank Harley Thompson for assisting with laboratory work (extracting DNA from cryptic species and helping with wing length assessments). For assistance in bee identification we thank Ken Walker at Museums Victoria and Michael Batley at the Australian Museum. We thank Australia Zoo for allowing us to sample on the Steve Irwin Wildlife Reserve in Cape York, Rawnsley Park Station in the Flinders Ranges, The Cairns Botanical Gardens, Big4 Carnarvon Gorge Caravan Park, Flinders University, The University of Queensland, and Monash University. We thank Nick Volpe for the use of native bee photos. Finally, we thank Michael Schwarz and Mark Stevens for early advice when planning the study.

## Author contributions

da Silva, Gloag and Kellermann conceptualised the study. da Silva led the field work and experiments with assistance from Beaman and Dorey. Bradford conducted the DNA extractions and library preparation for the ultraconserved element phylogeny. Smith led the morphological bee identification with assistance from da Silva and Dorey. da Silva collected wing length data. Kellermann and Gloag extracted DNA from cryptic species and species with high sample sizes that were not included in the UCE sequencing. Dorey constructed the multispecies ultraconserved element phylogeny. da Silva and Kellermann constructed the UCE – CO1 combined phylogeny. da Silva conducted the phylogenetic trait analyses. da Silva, Gloag and Kellermann wrote the first draft of the manuscript. All authors edited the manuscript.

## Data availability statement

The physiological trait data and climate data has been uploaded as a supplementary dataset and will be uploaded to a publicly available repository upon manuscript acceptance. The Australian native bee phylogeny has been uploaded as a supplementary file. The CO1 sequence data is available in GenBank (SUB15583452), and ultraconserved element sequence data is available in the NCBI Sequence Read Archive (PRJNA1314398).

