## Supplementary Tables and Figures for "Nesting behaviour predicts heat tolerance evolution and climate vulnerability in bees"

### Supplementary Material

*Supplementary Tables*

**S. Table 1.** Table of Australian native bee species and their mean collection latitudes, longitudes, heat tolerances, associated mean macro- and micro-climate data, and nesting behaviours. Bees were assigned nesting categories (ground, stem, or cavity nesting) based on nest type descriptions in taxonomic and reference texts (cited in Supplementary data). The category stem includes bees living in plant stems, twigs, and small narrow branches as they have similar thermal properties. Some species, such as those in the genus *Hylaeus*, can sometimes be found nesting in stems, twigs, or existing cavities in larger branches or trunks. For these species we allocated nesting strategies to the category where there were more numerous reports in the literature. Ultraconserved element sequence data is available in the NCBI Sequence Read Archive (PRJNA1314398). CO1 sequence data from the species we sequenced are available in GenBank (SUB15583452). CO1 data accession numbers from the literature (BOLD systems https://boldsystems.org/ and GenBank) are listed in the table below.

| Species | N samples | N sites | Heat tolerance | BIO5 | BIO12 | Microclimate temperature | Microclimate moisture | Nest | Bold Accession # |
|---|---|---|---|---|---|---|---|---|---|
| *Austroplebeia cassiae* | 47 | 5 | 41.30638 | 32.02 | 1287.15 | 28.29743 | 73.432 | Cavity | |
| *Austroplebeia australis* | 69 | 5 | 42.18261 | 32.84 | 906.9667 | 28.7073 | 69.24514 | Cavity | |
| *Tetragonula carbonaria* | 181 | 9 | 41.56022 | 30.38889 | 1248.5741 | 26.87234 | 72.71808 | Cavity | |
| *Tetragonula hockingsi* | 396 | 11 | 42.87753 | 31.55455 | 1295.4091 | 28.04086 | 72.32193 | Cavity | |
| *Tetragonula sapiens* | 24 | 1 | 43.22917 | 31.5 | 2832 | 28.2719 | 75.18405 | Cavity | |
| *Tetragonula clypearis* | 70 | 2 | 42.87571 | 32.65 | 2156 | 29.21476 | 76.63813 | Cavity | |
| *Braunsapis nitida* | 21 | 1 | 45.94762 | 31.5 | 2861.5 | 31 | 66.91459 | Stem | MSAPB2678-19 |
| *Braunsapis hyalina* | 6 | 2 | 44.65 | 31.25 | 1951.4166 | 30.7 | 63.73406 | Stem | |
| *Braunsapis unicolor* | 12 | 4 | 46.24167 | 31.6 | 1576.375 | 31.15 | 66.44498 | Stem | |
| *Braunsapis protuberans* | 43 | 6 | 46.14884 | 30.31667 | 1675.6111 | 29.9 | 65.79287 | Stem | OZBOL157-21 |
| *Braunsapis semillina* | 5 | 4 | 46.52 | 32.025 | 1355.7083 | 31.9 | 62.68193 | Stem | MSAPB2674-19 |

| | | | | | | | | | |
|---|---|---|---|---|---|---|---|---|---|
| *Braunsapis plebeia* | 11 | 2 | 46.87273 | 32.5 | 1312.4166 | 31.8 | 65.6728 | Stem | |
| *Braunsapis clarissima* | 7 | 4 | 44.6 | 31.625 | 1753.25 | 31.15 | 66.44498 | Stem | |
| *Exoneura angophorae* | 10 | 3 | 42.59 | 29.13333 | 1684.5 | 28.6 | 64.34172 | Stem | |
| *Brevineura Adelaide morph 1* | 11 | 1 | 45.47273 | 27.7 | 596.2 | 27.3 | 57.98156 | Stem | |
| *Megachile micrerythrura* | 5 | 2 | 43.76 | 32.85 | 1113.75 | 29.13826 | 71.22045 | Cavity | |
| *Megachile punctata* | 6 | 1 | 47.03333 | 31.7 | 644 | 28.11887 | 64.3487 | Cavity | |
| *Megachile deanii* | 45 | 11 | 44.9 | 31.14545 | 1174.9864 | 27.43846 | 71.4206 | Cavity | |
| *Megachile abdominale* | 6 | 2 | 43.68333 | 32.5 | 626.5 | 28.14098 | 66.03112 | Cavity | MSAPB3161-19 |
| *Xanthesma brachycera* | 9 | 2 | 46.56667 | 32.5 | 626.5 | 23.08196 | 99.99215 | Ground | |
| *Callohesma calliopsiformis* | 43 | 2 | 39.90233 | 29.5 | 893.4 | 22.02631 | 99.99219 | Ground | |
| *Euryglossa morpho species 1* | 6 | 1 | 45.53333 | 30 | 1526.6667 | 23.10069 | 99.99226 | Ground | |
| *Euryglossa ephippiata* | 18 | 1 | 42.4 | 26.8 | 836.75 | 17.98226 | 99.99204 | Ground | MSAPB3241-19 |
| *Euryglossa adelaidae* | 4 | 2 | 45.125 | 27.25 | 717.175 | 18.30651 | 99.99206 | Ground | PMSAPB3178-19 |
| *Hyphesma atromicans* | 4 | 2 | 42.925 | 29.7 | 620.8 | 20.93425 | 99.99213 | Ground | MSAPB1438-19 |
| *Euryglossina morpho species 1* | 8 | 1 | 45.7375 | 27.7 | 597.6 | 27.3 | 57.98156 | Stem | |
| *Euryglossina mellea* | 6 | 1 | 45.73333 | 26.8 | 836.75 | 25.9 | 62.90322 | Stem | OZBOL043-21 |
| *Euryglossina lynettae* | 4 | 1 | 44.475 | 27.6 | 1171 | 27 | 63.24236 | Stem | MSAPB3883-19 |
| *Euryglossina morpho species 2* | 44 | 1 | 41.62727 | 31 | 1041.3333 | 30.4 | 60.55353 | Stem | |
| *Euryglossina hypochroma* | 86 | 3 | 44.83953 | 29.23333 | 701.25 | 28.86667 | 59.2796 | Stem | |

| | | | | | | | | | |
|---|---|---|---|---|---|---|---|---|---|
| *Euryglossula fultoni* | 20 | 2 | 43.33 | 30.35 | 495 | 20.00189 | 99.99191 | Ground | |
| *Pachyprosopis kellyi* | 86 | 4 | 42.08605 | 28.775 | 797.9 | 28.4 | 60.2712 | Stem | |
| *Pachyprosopis haematostoma* | 31 | 4 | 44.23548 | 28.775 | 792.5875 | 24.27471 | 70.20075 | Cavity | OZBOL2714-21 |
| *Pachyprosopis holoxanthopus* | 4 | 2 | 43.825 | 30.5 | 603.3 | 30.35 | 57.46779 | Stem | AUSBS374-13 |
| *Pachyprosopis xanthodonta* | 39 | 1 | 43.6 | 33.3 | 609 | 33.4 | 56.95402 | Stem | |
| *Pachyprosopis mirabillis* | 6 | 3 | 45.96667 | 32.83333 | 769.1111 | 32.83333 | 57.2403 | Stem | |
| *Hylaeus albonitens* | 249 | 10 | 45.0988 | 30.82 | 1448.6267 | 30.45 | 64.73444 | Stem | AUSBS300-13 |
| *Hylaeus euxanthus* | 88 | 10 | 45.49205 | 29.8 | 1018.4783 | 29.49 | 59.90074 | Stem | GCQT846-17 |
| *Hylaeus chromaticus* | 4 | 2 | 41.375 | 30.65 | 1146.5 | 29.8 | 60.60889 | Stem | |
| *Hylaeus philoleucus* | 34 | 1 | 45.52647 | 27.6 | 1171 | 25.9 | 62.90322 | Stem | |
| *Hylaeus eugeniellus* | 15 | 5 | 42.05333 | 28.46 | 1009.9167 | 27.96 | 61.06899 | Stem | |
| *Hylaeus minusculus* | 4 | 1 | 44.325 | 26.8 | 836.75 | 25.9 | 62.90322 | Stem | |
| *Hylaeus morpho species 3* | 5 | 1 | 44.14 | 28.4 | 1728.5 | 27.8 | 62.86819 | Stem | |
| *Hylaeus littleri* | 6 | 2 | 44.25 | 27 | 981.675 | 26.45 | 63.07463 | Stem | AUSMD422-14 |
| *Hylaeus aralis* | 146 | 8 | 43.76575 | 29.5625 | 882.8854 | 29.225 | 59.92695 | Stem | MSAPB1485-19 |
| *Hylaeus perhumilis* | 19 | 3 | 43.62105 | 27.23333 | 900.5167 | 26.73333 | 61.37571 | Stem | MSAPB1514-19 |
| *Hylaeus chlorosoma* | 140 | 7 | 42.89714 | 31.08571 | 684.1976 | 21.89197 | 99.9921 | Ground | MSAPB1505-19 |
| *Hylaeus morpho species1* | 10 | 1 | 43.28 | 28.4 | 1728.5 | 27.8 | 62.86819 | Stem | |
| *Hylaeus microphenax* | 10 | 2 | 43.42 | 30.65 | 1160.9166 | 29.8 | 60.60889 | Stem | |

| | | | | | | | | | |
|---|---|---|---|---|---|---|---|---|---|
| *Hylaeus primulipictus* | 8 | 3 | 43.8125 | 28.1 | 1369.1778 | 28 | 65.62493 | Stem | |
| *Hylaeus elegans* | 35 | 5 | 42.13143 | 31.04 | 851.5533 | 22.29048 | 99.99212 | Ground | AUSBS380-13 |
| *Hylaeus disjunctus* | 4 | 3 | 43.525 | 30.43333 | 1245.6667 | 29.86667 | 62.91251 | Stem | |
| *Hylaeus lateralis* | 24 | 5 | 45.00417 | 30.44 | 962.5567 | 30.04 | 61.83689 | Stem | OZBOL2729-21 |
| *Hylaeus rotundiceps* | 5 | 3 | 42.44 | 27.46667 | 1230.6167 | 23.66015 | 72.2524 | Cavity | MSAPB3037-19 |
| *Hylaeus ruficeps ruficeps* | 22 | 7 | 44.35 | 31.7 | 1115 | 27.77587 | 69.93842 | Cavity | MSAPB1508-19 |
| *Hylaeus cyanurus* | 4 | 1 | 41.425 | 31.4 | 2420.6667 | 28.27187 | 75.18405 | Cavity | |
| *Hylaeus violaceus* | 5 | 1 | 45.32 | 30.3 | 1169 | 29.3 | 60.82062 | Stem | MSAPB1683-19 |
| *Hylaeus nubilosus* | 12 | 5 | 41.36667 | 27.82 | 1053.0967 | 27.28 | 62.18082 | Stem | |
| *Meroglossa torrida* | 15 | 1 | 45.09333 | 34.2 | 661 | 34.7 | 54.21335 | Stem | MSAPB4791-19 |
| *Palaeorhiza parallela* | 11 | 3 | 40.8 | 30.6 | 1852.3889 | 23.93486 | 99.99231 | Ground | HYQT617-10 |
| *Palaeorhiza melanura* | 4 | 1 | 40.8 | 31.5 | 2861.5 | 28.27189 | 75.18404 | Cavity | |
| *Leioproctus morpho species 1* | 4 | 1 | 42.85 | 27.6 | 1171 | 21.05888 | 99.99217 | Ground | |
| *Leioproctus clarki* | 8 | 2 | 40.1875 | 27.2 | 747.375 | 18.30652 | 99.99206 | Ground | MSAPB1582-19 |
| *Leioproctus amabilis* | 5 | 2 | 39.84 | 30.45 | 867.5 | 21.99242 | 99.99213 | Ground | |
| *Leioproctus carinatus* | 14 | 2 | 38.60714 | 29.9 | 595.25 | 19.67767 | 99.99197 | Ground | |
| *Leioproctus launcestonensis* | 43 | 1 | 41.45349 | 26.8 | 836.75 | 17.98226 | 99.99204 | Ground | |
| *Ceylalictus perditellus* | 51 | 4 | 44.31765 | 30.65 | 1394.75 | 23.39596 | 99.99214 | Ground | |
| *Homalictus eurhodopus* | 29 | 2 | 41.24483 | 30.75 | 2194.0833 | 24.32225 | 99.99234 | Ground | |
| *Homalictus blackburni* | 22 | 3 | 41.73636 | 30.83333 | 1809.8333 | 23.99759 | 99.99228 | Ground | GCQT1058- |

| | | | | | | | | | |
|---|---|---|---|---|---|---|---|---|---|
| | | | | | | | | | 17 |
| *Homalictus murrayi* | 8 | 2 | 43.725 | 30.85 | 1085.3334 | 23.16922 | 99.99223 | Ground | MSAPB1224-12 |
| *Homalictus flindersi* | 63 | 3 | 42.53175 | 29.8 | 1432.1667 | 22.97097 | 99.99222 | Ground | BOWAU016-10 |
| *Homalictus cassiaefloris* | 26 | 3 | 42.98462 | 30.83333 | 1809.8333 | 23.99759 | 99.99228 | Ground | |
| *Homalictus stradbrokensis* | 19 | 2 | 41.61579 | 30.65 | 1167.4166 | 23.39603 | 99.9922 | Ground | |
| *Homalictus dampieri* | 21 | 5 | 43.05238 | 30.98 | 1695.6 | 24.31441 | 99.99233 | Ground | AUSBC1624-12 |
| *Homalictus urbanus* | 20 | 5 | 42.26 | 31.44 | 1027.02 | 23.65673 | 99.99227 | Ground | ASMII7111-22 |
| *Homalictus megastigmus* | 7 | 2 | 40.2 | 26.2 | 915.5417 | 17.84357 | 99.99205 | Ground | |
| *Homalictus brisbanensis* | 45 | 3 | 41.39111 | 28.16667 | 1086.1389 | 20.74707 | 99.99216 | Ground | |
| *Homalictus punctatus* | 88 | 3 | 40.92045 | 27.06667 | 896.45 | 19.14255 | 99.9921 | Ground | |
| *Homalictus sphecodopsis* | 17 | 4 | 41.91765 | 28.15 | 1455.6083 | 22.03625 | 99.99214 | Ground | MSAPB1207-1 |
| *Homalictus dotatus* | 83 | 7 | 42.69518 | 31.18571 | 857.0262 | 22.52717 | 99.99215 | Ground | |
| *Homalictus caloundrensis* | 113 | 5 | 41.67611 | 30.24 | 1483.5 | 23.20957 | 99.99216 | Ground | |
| *Homalictus behri* | 72 | 1 | 42.59167 | 31.5 | 2598 | 25.54381 | 99.99243 | Ground | |
| *Lasioglossum bicingulatum* | 141 | 5 | 42.70426 | 27.44 | 1147.3533 | 20.33154 | 99.99208 | Ground | |
| *Lasioglossum morpho species 3* | 18 | 1 | 41.90556 | 26.8 | 836.75 | 17.98226 | 99.99204 | Ground | |
| *Lasioglossum mediopolitum* | 14 | 1 | 43.30714 | 33 | 351.5 | 21.37307 | 99.99175 | Ground | MH320169* GenBank |
| *Lasioglossium gilesi* | 16 | 1 | 41.91818 | 27.6 | 658 | 18.63078 | 99.99207 | Ground | |
| *Lasioglossum hemichalceum* | 27 | 2 | 43.36296 | 27.2 | 747.375 | 18.30652 | 99.99206 | Ground | |

| | | | | | | | | | |
|---|---|---|---|---|---|---|---|---|---|
| *Lasioglossum appositum* | 8 | 1 | 44.9875 | 31.7 | 644 | 23.23774 | 99.9922 | Ground | |
| *Lasioglossum quadratum* | 6 | 1 | 45.48333 | 33.3 | 609 | 22.92618 | 99.99211 | Ground | OZBOL216-21 |
| *Lasioglossum vitripenne* | 15 | 2 | 42.85333 | 29.9 | 594.75 | 19.67767 | 99.99196 | Ground | OZBOL251-21 |
| *Lasioglossum sulthicum* | 10 | 3 | 41.65 | 27.13333 | 1150.25 | 19.69233 | 99.99212 | Ground | |
| *Lasioglossum hiltacum* | 26 | 3 | 41.96923 | 29.23333 | 688.6944 | 19.84641 | 99.99207 | Ground | |
| *Lasioglossum leichardti* | 9 | 2 | 41.22222 | 27.7 | 1584.5834 | 22.1148 | 99.99206 | Ground | |
| *Patellapis stirlingi* | 10 | 1 | 41.74 | 30 | 1526.6667 | 23.10069 | 99.99226 | Ground | |
| *Lipotriches flavoviridis* | 35 | 2 | 45.40857 | 28.6 | 1326.6334 | 22.02108 | 99.99217 | Ground | |

**S. Table 2**. Phylogenetic mixed model AICc comparison using the program *slouch* (Kopperud et al., 2024). Models were made using either an Ournsein-Uhlenbeck (OU) or Brownian Motion (BM) model of evolution. Wing length was included into the best climate predictor model. Whether wing length played a significant role in explaining variation in heat tolerance was tested by conducting a likelihood ratio test. Wing length did not explain significant variation in heat tolerance (LRT: $\chi^2$ = 2.01, $p$ = 0.156). Thus, we discuss the model parameters from the best fitting model without wing length despite there being an AICc difference of less than two. The similar model performance is due to microclimate temperature explaining most of the variation in the data.

| Model of evolution | Variables | AICc | $R^2$ | $t_{1/2}$ |
|---|---|---|---|---|
| **OU** | **micro temp + phy** | **349.87** | **0.1247** | **26.64** |
| OU | micro temp + wing length + phy | 350.1 | 0.145 | 25.44 |
| OU | micro temp + micro humidity + phy | 351.7724 | 0.1279 | 25.97 |
| OU | micro humidity + phy | 353.81 | 0.084 | 25.98 |
| BM | micro temp + phy | 353.89 | 0.099 | |
| OU | bio5 + phy | 355.4782 | 0.06055 | 37.75 |
| BM | micro temp + micro humidty + phy | 356 | 0.1 | |
| BM | bio5 + phy | 356.85 | 0.07 | |
| OU | bio5 + bio12 + phy | 357.0011 | 0.06804 | 38.78 |
| BM | bio5 + bio12 + phy | 357.57 | 0.085 | |
| OU | phy | 359 | 0 | 35.79 |
| OU | bio12 + phy | 360.6 | 0.0056 | 36.76 |
| BM | phy | 362 | -1.46E-16 | |

**S. Table 3.** Model parameter values from the microclimate maximum temperature phylogenetic mixed model (*slouch*).

| **Inferred maximum likelihood parameters** | **Parameter values** |
|---|---|
| Mean phylogenetic correction factor | 0.655 |
| Rate of adaptation | 0.026 |
| Diffusion variance | 0.112 |
| Optimal regression slope | 0.286 ± 0.078 |
| Evolutionary regression slope | 0.187 ± 0.051 |

**S Table 4.** Average number of nesting behavior transitions across the phylogenetic tree based on 1,000 trees with ancestral nesting behaviour mapped across the branches (trees have 17.22 changes between nesting states on average).

| **Nesting behaviour transition** | **Number of transitions** |
|---|---|
| Cavity to ground | 1.623 |
| Cavity to stem | 1.855 |
| Ground to cavity | 2.022 |
| Ground to stem | 2.352 |
| Stem to cavity | 3.898 |
| Stem to ground | 5.47 |

**S. Table 5.** Comparison of models to determine how nesting strategy shapes heat tolerance evolution. BM = Brownian Motion; OU = Ornstein Ulenbeck. BM1 and OU1 models do not depend on internal nesting states and therefore there was no need to conduct 1000 simulations and calculate confidence intervals. We had insufficient data to test a model that included multiple evolutionary rates and optima depending on nesting behaviour, this over parameterised model was unreliable and was not included in the comparison. Using such multi-rate and optima models is generally not recommended (Revell & Harmon, 2022).

| **Model** | **Parameters** | **2.5 CI AICc** | **Mean AICc** | **97.5 CI AICc** |
|---|---|---|---|---|
| OUM | Single rate, multi optima | 362.72 | **358.37** | 366.61 |
| OU1 | Single rate, single optima | | 365.28 | |
| BM1 | single rate, no optima | | 381.62 | |
| BMS | Multi rate, no optima | 383.25 | 381.31 | 384.36 |

**Table 6.** Multi-regime OU heat tolerance evolution model parameter outputs (mean and 95% confidence intervals) from 1000 nesting ecology ancestral state simulations. $\alpha$ = strength of selection on heat tolerance evolution, $\sigma^2$ = rate of heat tolerance evolution (°C per million years), $\boldsymbol{\theta}$ = regieme mean (°C). Note that the evolutionary rate parameter $\sigma^2$ should not be interpreted as the maximum rate heat tolerance can evolve as it is averaged across the entire phylogeny (104 MY).

| Parameter | Mean ($\mu$) | Low 95% CI | High 95% CI |
|---|---|---|---|
| $\alpha$ | 5.94 | 4.72 | 6.79 |
| $\sigma^2$ | 0.34 | 0.30 | 0.36 |
| Stem $\boldsymbol{\theta}$ | 44.90 | 44.06 | 45.43 |
| Cavity $\boldsymbol{\theta}$ | 43.27 | 42.34 | 43.93 |
| Ground $\boldsymbol{\theta}$ | 42.88 | 42.32 | 43.45 |

**S. Table 7.** The number of individuals assessed for heat tolerance each month for each of the ten species used in the analysis to partition variation in heat tolerance to the species and month.

| Species | month | count |
|---|---|---|
| *Euryglossina hypchroma* | Nov | 3 |
| | Feb | 27 |
| | March | 43 |
| *Euryglossina mellea* | Feb | 3 |
| | March | 3 |
| *Homalictus brisbanensis* | Jan | 5 |
| | Feb | 12 |
| | March | 9 |
| *Hylaeus aralis* | Jan | 9 |
| | Feb | 53 |
| | March | 14 |
| *Hylaeus euxanthus* | Nov | 11 |
| | Jan | 11 |
| | Feb | 4 |
| *Hylaeus philolecus* | Jan | 4 |
| | Feb | 26 |
| *Lasioglossium bicingulatum* | Nov | 6 |
| | Jan | 52 |
| | Feb | 5 |
| | Mar | 15 |
| *Lasioglossium hemichalceum* | Jan | 3 |
| | Feb | 5 |
| *Pachprosopis kellyi* | Feb | 18 |
| | Mar | 11 |

**S Table 8.** Field trip order and locations. Two field trips were conducted to collect the native bee heat tolerance dataset. To account for the effect of season during the field trips on bee heat tolerance, the first field trip was conducted predominantly in a north to south direction. The first part of the second field trip was conducted in a north to south direction, and the second half was collected predominantly north to south across eastern Australia (inland and coastal sites). GPS coordinates for each location are in brackets.

| **Field trip 1 collection order** | **Field trip 2 collection order** |
|---|---|
| Melbourne VIC (-37.9143, 145.1291) | Adelaide SA (-35.0307, 138.5788) |
| Adelaide SA (-35.0307, 138.5788) | Flinders Ranges SA (-31.6443, 138.5800) |
| Melbourne VIC (-37.9143, 145.1291) | Pentland QLD (-20.5247, 145.4017) |
| Brisbane QLD (-27.5008, 153.0163) | Cairns QLD (-16.8867, 145.7486) |
| Cairns QLD (-16.8867, 145.7486) | Steve Irwin Wildlife Reserve (-12.3741, 142.1959) |
| Sydney NSW (-33.9023, 151.1616) | Melbourne VIC (-37.9143, 145.1291) |
| Adelaide SA (-35.0279, 138.5731) | Townsville QLD (-19.3749, 146.7310) |
| Melbourne VIC (-37.9143, 145.1291) | Finch Hatton QLD (-21.1363, 148.5106) |
| | Sarina QLD (-21.4262, 149.2182) |
| | Melbourne VIC (-37.9143, 145.1291) |
| | Yepoon QLD (-23.1220, 150.7320) |
| | Carnarvon Gorge QLD (-25.0715, 148.2709) |
| | Melbourne VIC (-37.9143, 145.1291) |
| | Miriam Vale QLD (-24.3278, 151.5640) |
| | Ocean Shores NSW (-28.3878, 153.5661) |
| | Pilliga NSW (-30.9459, 149.0686) |
| | South West Rocks NSW (-30.8878, 153.0405) |
| | Sydney NSW (-33.7606, 151.1021) |

**S Table 9.** Heat tolerance evolution hypothesis comparison table. Local climates = maximum temperature of the hottest month (BIO5) and annual precipitation (BIO12). Microclimate temperature = hottest temperature within the nest (mean across collection locations) (ground, cavity, or stem). Nest factor = ground, cavity or stem.

| **Hypothesis** | **Variables** |
|---|---|
| Only phylogeny explains $CT_{MAX}$ evolution | Phylogeny |
| Broadscale climate predicts $CT_{MAX}$ evolution (controlling for phylogenetic inertia) | Maximum temperature + annual precipitation + phylogeny |
| Broadscale temperature predicts $CT_{MAX}$ evolution (controlling for phylogenetic inertia) | Maximum temperature + phylogeny |
| Broadscale precipitation predicts $CT_{MAX}$ evolution (controlling for phylogenetic inertia) | Maximum precipitation + phylogeny |
| Nesting microclimates explains $CT_{MAX}$ evolution (controlling for phylogenetic inertia) | Microclimate temperature + microclimate humidity + phylogeny |
| Microclimate temperature explains $CT_{MAX}$ evolution (controlling for phylogenetic inertia) | Microclimate temperature + phylogeny |
| Microclimate humidity explains $CT_{MAX}$ evolution (controlling for phylogenetic inertia) | Microclimate humidity + phylogeny |
| Wing length as a proxy for body mass was added into the best fitting model | Microclimate temperature + wing length + phylogeny |

**Supplementary Figures**

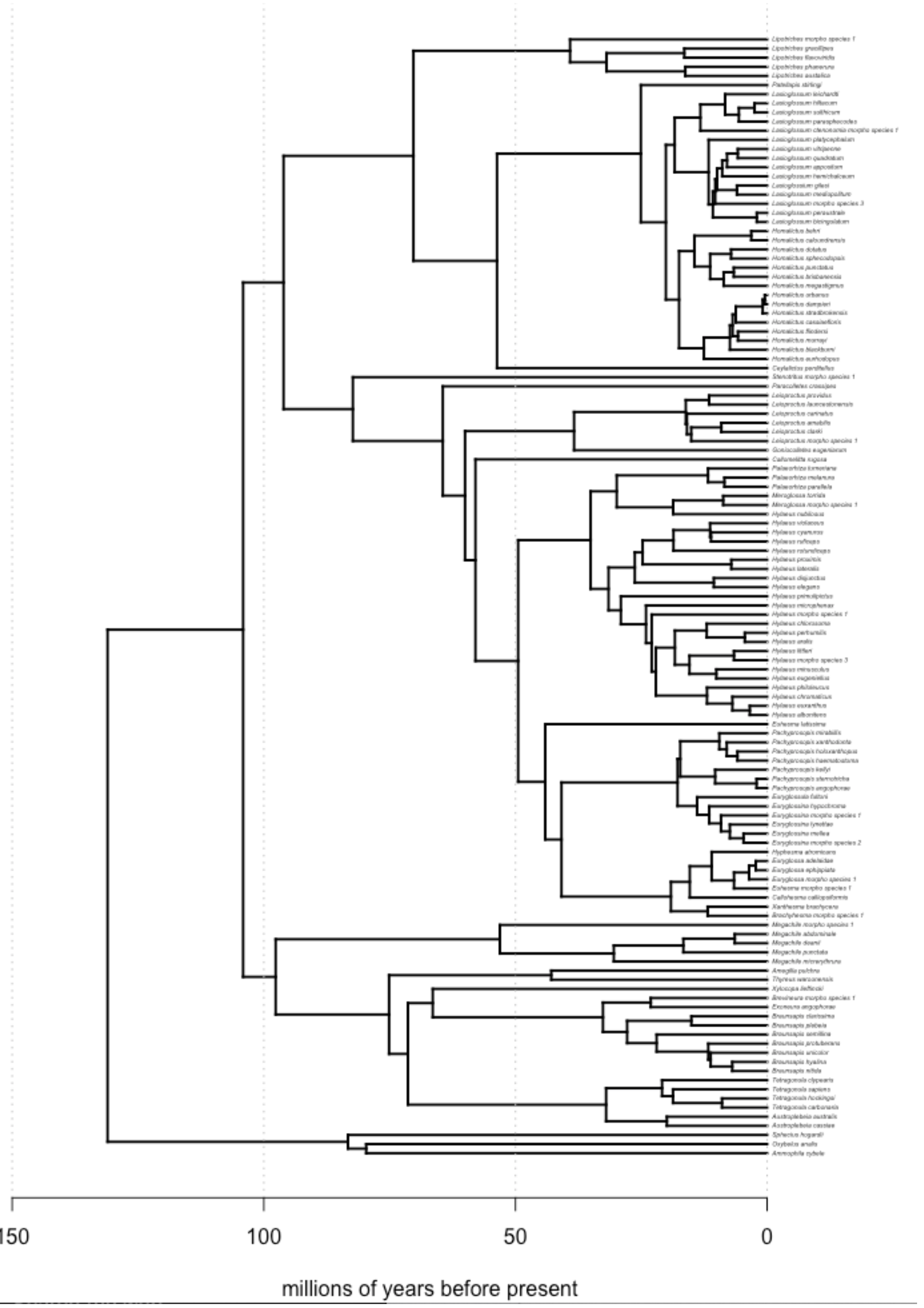

**S Figure 1.** Time calibrated Australian native bee phylogeny. Note that more species of bee are included in the phylogeny than the dataset. This is because some species did not have high enough sample sizes to be included in the phylogeny. However, for interest in bee phylogenetics and to have as full of a phylogeny as possible, they were included in the phylogeny but pruned out for the phylogenetic trait analysis. In this phylogeny, there are 78 UCE samples which make up the backbone of the phylogeny, and 45 CO1 samples (N = 123). The final phylogeny used in our analysis included 52 UCE samples and 43 CO1 samples (N = 95).

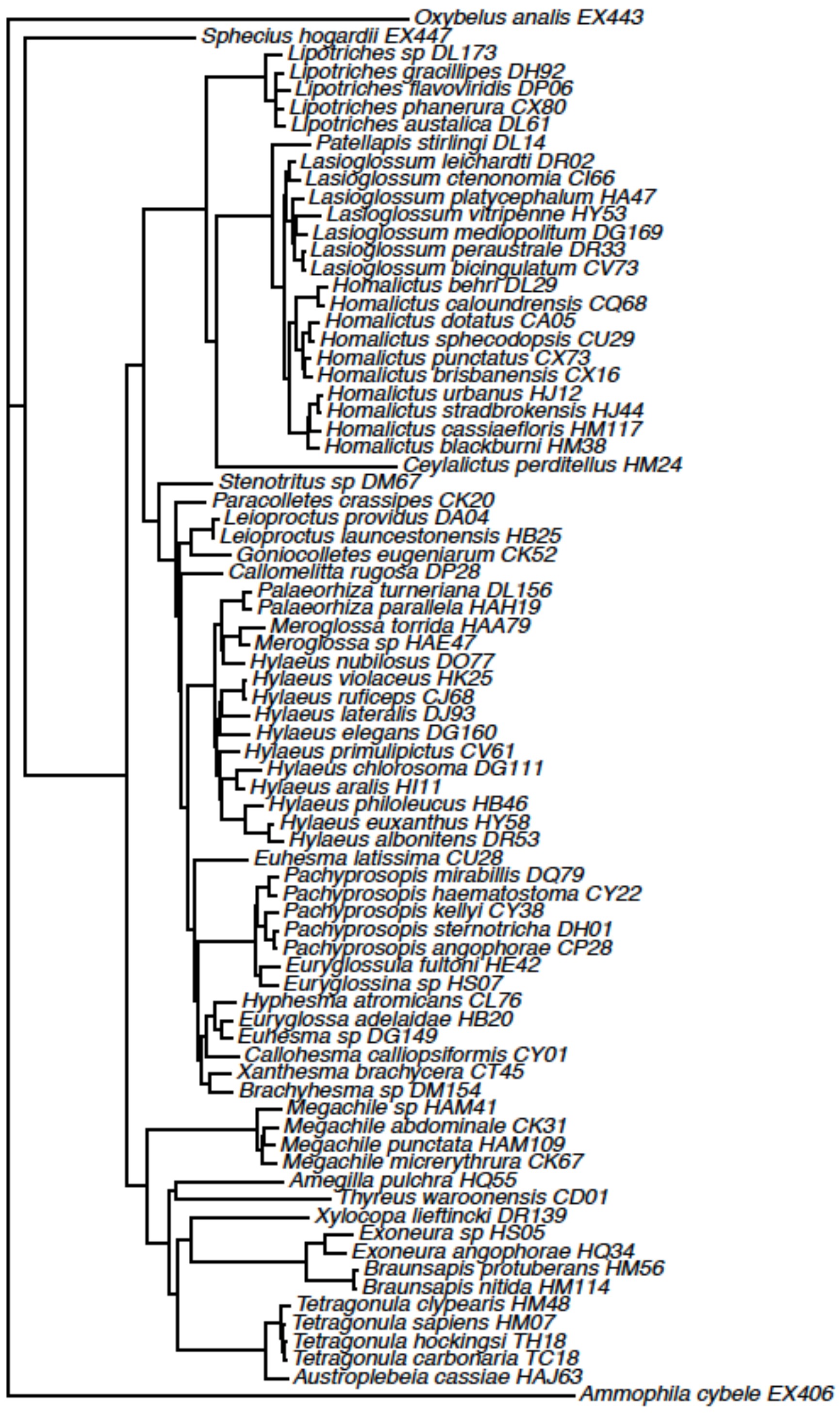

**S Figure 2**. Species included in the ultra-conserved element phylogeny.

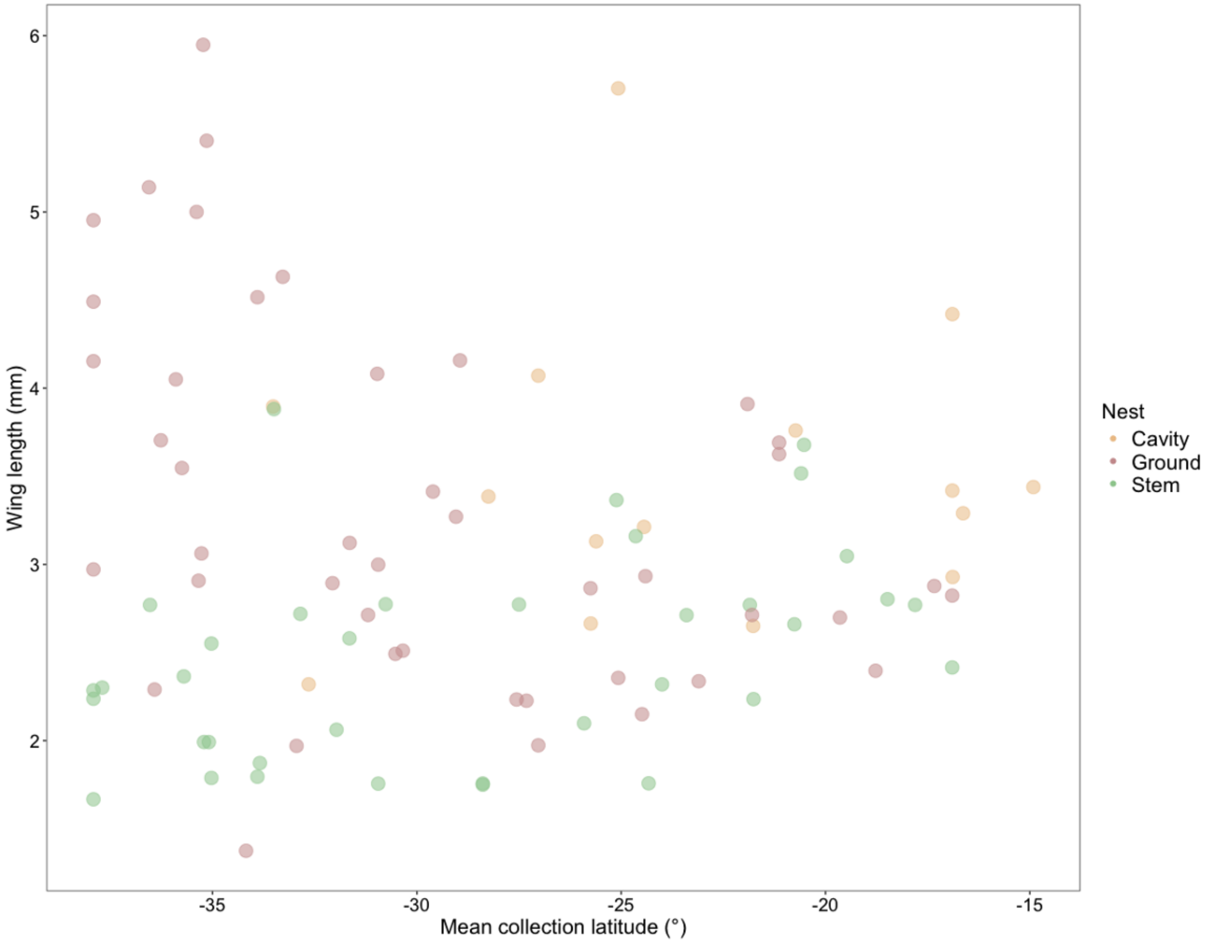

**S Figure 3.** No association between wing length (mm) and mean collection latitude (LRT: $\chi^2$ = 1.85, $p$ = 0.174). To ensure that the large high latitude bees in this dataset were not obscuring any signal in wing length patterns across latitude, we re-ran the analysis excluding bees that had wing lengths over 4mm. However, wing length, as a proxy for body size, still did not change predictably with latitude (LRT: $\chi^2$ = 327, $p$ = 0.567).

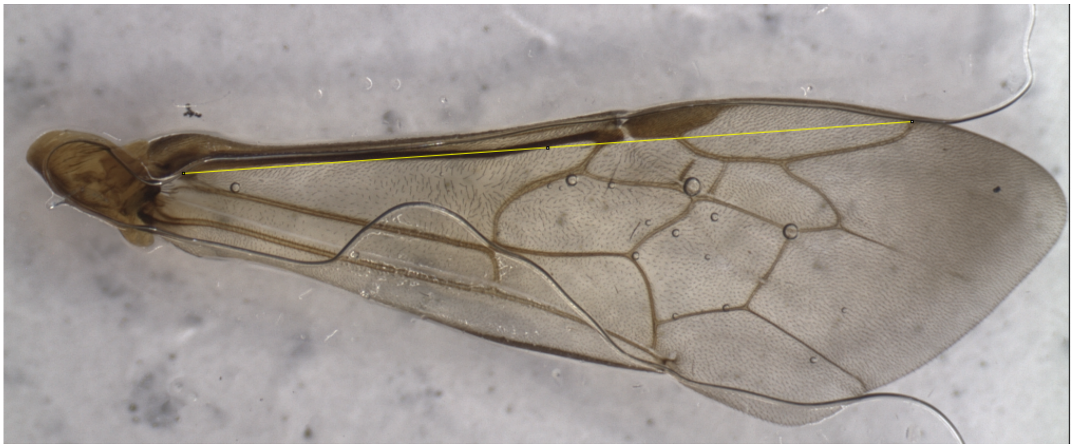

**S Figure 4.** Example of how wing length was measured.

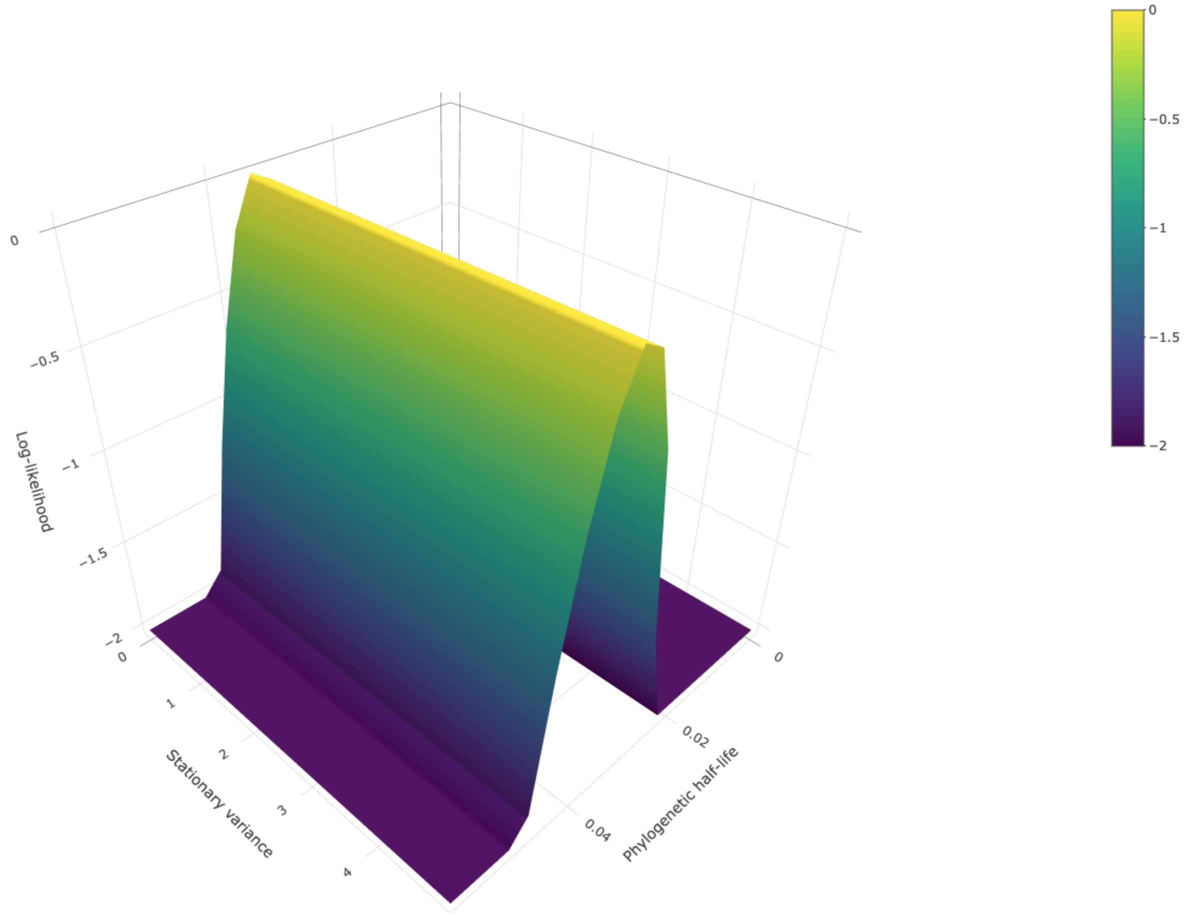

**S Figure 5.** Phylogenetic inertia in the heat tolerance of Australian native bees. Grid search support surfaces for the intercept (phylogeny) only model phylogenetic half-life ($t_{1/2}$), stationary variance, and support (log-likelihood).

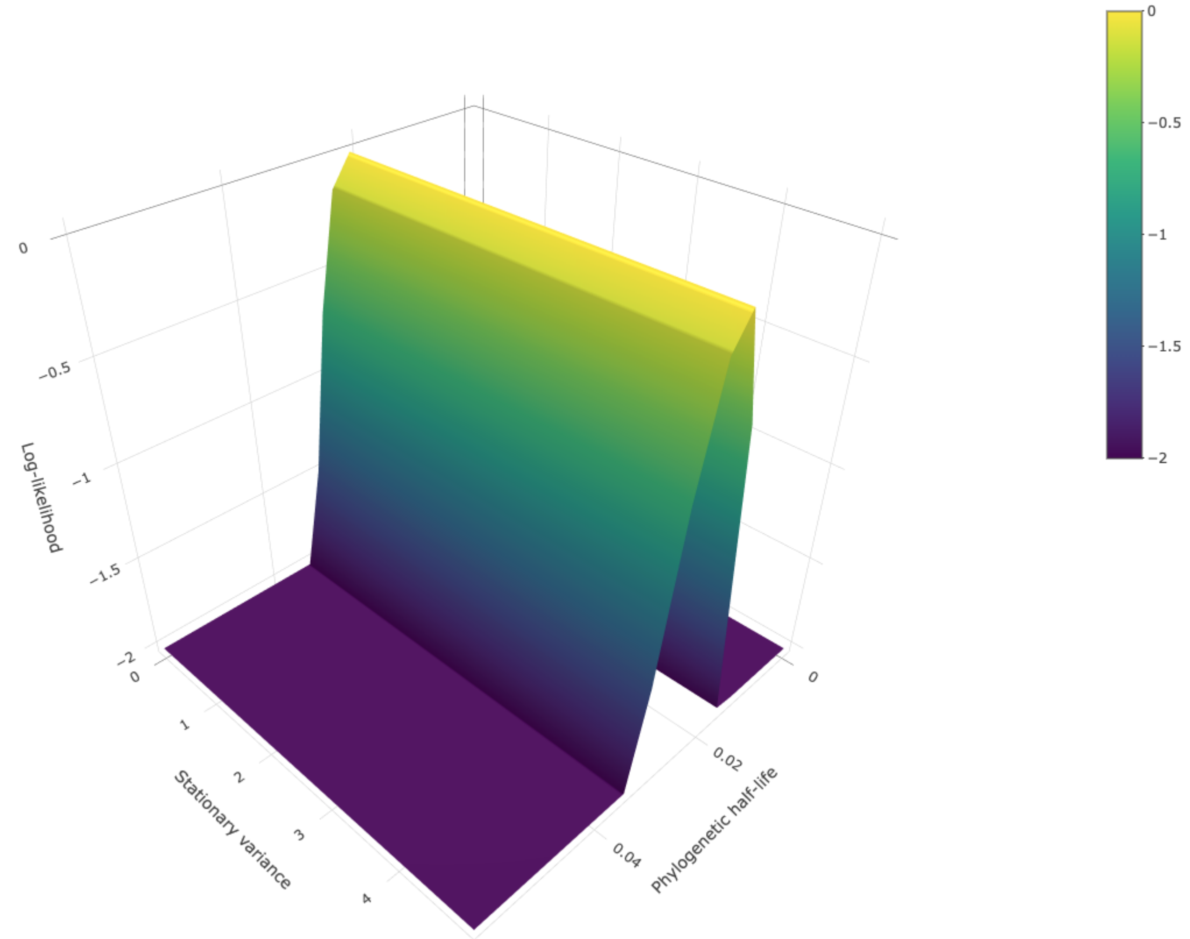

**S Figure 6**. Phylogenetic effects the heat tolerance of Australian native bees. Grid search support surfaces for the microclimate temperature model phylogenetic half-life ($t_{1/2}$), stationary variance, and support (log-likelihood).

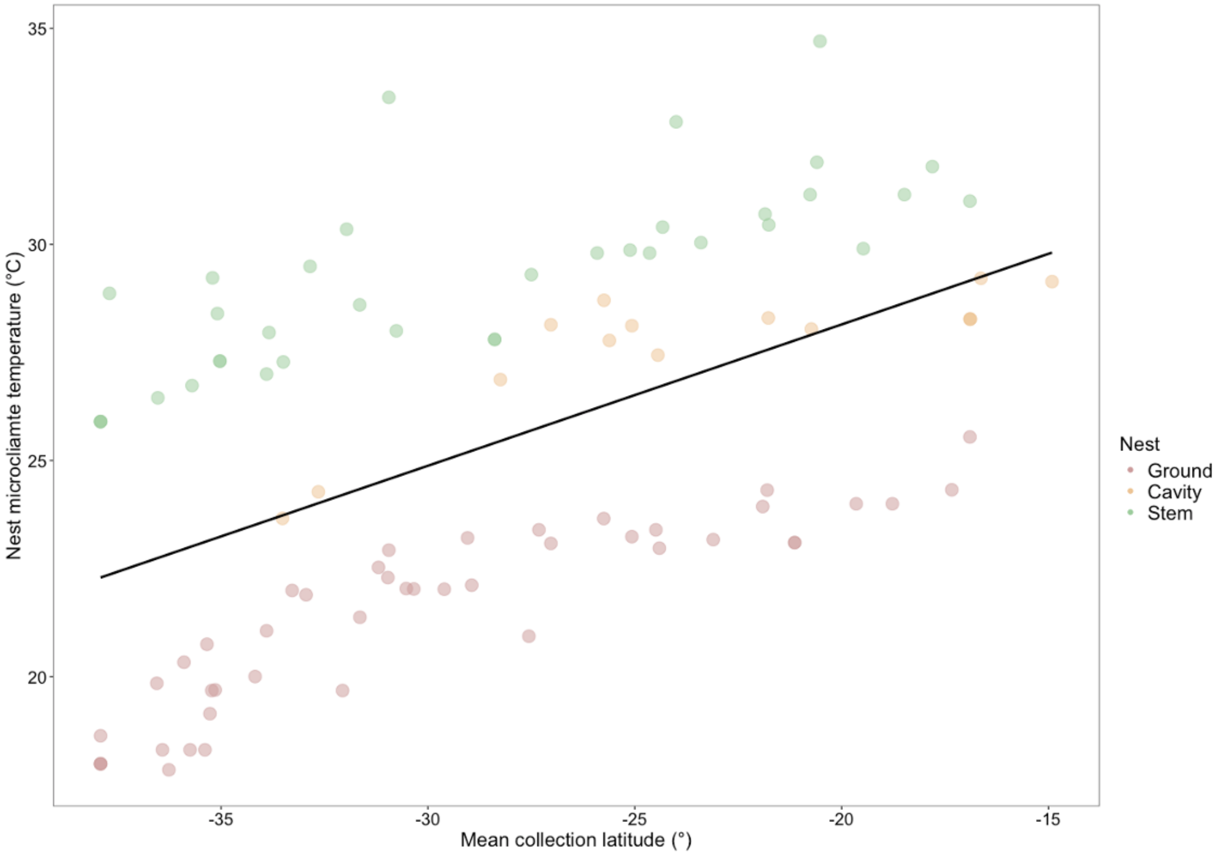

**S Figure 7.** Relationship between maximum nest microclimate temperature and latitude ($R^2$ = 0.54).
